# Probabilistic inference of Homonymous and Heteronymous Recurrent Inhibition in Human Muscles from Large-Scale Motor Neuron Recordings

**DOI:** 10.64898/2025.12.19.694963

**Authors:** François Dernoncourt, Simon Avrillon, Thomas Cattagni, Dario Farina, François Hug

**Affiliations:** Université Côte d’Azur, LAMHESS, Nice, France; Department of Bioengineering, Faculty of Engineering, Imperial College London, London, United Kingdom; Nantes Université, Movement-Interactions-Performance, MIP, UR 4334, F-44000, Nantes, France; The University of Queensland, School of Biomedical Sciences, Brisbane, QLD, Australia; Institut Universitaire de France (IUF), Paris, France

## Abstract

Understanding how spinal circuits shape motor neuron behavior during muscle contractions remains a major challenge. Here, we combined large-scale motor unit recordings with simulation-based inference to generate probabilistic estimates of homonymous and heteronymous recurrent inhibition, a key spinal circuit that has remained largely inaccessible during natural voluntary contractions. We constructed synchronization cross-histograms from motor neuron spike trains and extracted features representative of recurrent inhibition. Because these features are also influenced by higher-frequency components of common synaptic input, we developed a simulation-based inference framework to disentangle these effects. Following validation, we applied this framework to experimental data from six muscles at two contraction intensities, revealing previously uncharacterized muscle-and intensity-dependent patterns: recurrent inhibition decreased with contraction intensity in most muscles but increased in the vastus lateralis and medialis. The pipeline is openly available and designed for reuse on comparable datasets and for adaptation to diverse experimental contexts, including other spinal circuits.

**Teaser:** Large-scale motor-unit recordings enable in-vivo probabilistic mapping of spinal recurrent inhibition.

## Introduction

Understanding how synaptic inputs shape motor neuron activity and ultimately generate muscle contraction remains limited. At the population level, motor neuron activity dynamics emerge from the interplay of descending supraspinal drive, sensory feedback, and spinal circuits, and exhibit substantial variation across motor neuron pools (*1–3*). Yet, inferring the contribution of specific spinal circuits to motor neuron behavior remains a central barrier to understanding the neural control of movement.

Among spinal circuits, recurrent inhibition is a key mechanism that shapes motor neuron dynamics. Motor axon collaterals excite Renshaw cells that, in turn, inhibit the motor neurons they contact, forming a disynaptic negative feedback loop (*4*, *5*). Despite these effects of recurrent inhibition on motor neuron behavior, its functional role in motor execution remains poorly understood. Proposed roles of recurrent inhibition include tremor-band damping through frequency-selective filtering (*6*, *7*), as well as contributing to error correction and motor adaptation/learning (*8–10*). These roles imply muscle-and task-dependent expression of recurrent inhibition. For example, because limb mechanical resonance frequency is generally higher in distal than in proximal joints (*11*), a recurrent loop that attenuates resonant oscillations at proximal joints could amplify them at distal joints, consistent with evidence of weaker recurrent inhibition in distal motor neuron pools (*6*). Clarifying the role of recurrent inhibition therefore requires accurate characterization across motor neuron pools and contraction intensities.

Recurrent inhibition has been quantified in reduced or anesthetized animal preparations (*6*, *12*). Although these studies provide precise measurements, their results cannot be directly extrapolated to natural behavior. In humans, mapping the distribution of recurrent inhibition across muscles has been attempted using indirect stimulation-based approaches such as the paired H-reflex and M-wave (*13–15*). These methods have been invaluable for probing recurrent inhibition, but they suffer from substantial limitations. First, stimulation imposes non-physiological patterns of motor neuron recruitment, producing responses that may not reflect spinal circuit behavior during voluntary contractions. Second, these methods sample a biased subset of motor neurons. Specifically, the paired H-reflex method assesses recurrent inhibition only in the subset of motor neurons recruited by the stimulation-induced Ia afferent volley (*16*), whereas the M-only paradigm preferentially measures inhibition directed from large to small motor neurons (*17*). Third, the paired H-reflex method is susceptible to confounds from intrinsic motor neuron properties because it recruits the same motor neurons twice within a short interval (*18*). Finally, these approaches can be used only in muscles that are easy to stimulate. As a result, human data are largely confined to the triceps surae, leaving many other pools insufficiently characterized.

Approaches have recently been developed to reverse-engineer the synaptic inputs received by motor neurons from their output, i.e., their firing patterns (*19*). The underlying principle is that, with a physiologically grounded model, motor neuron spiking activity can be simulated under different input scenarios to determine which ones best match the experimental observations. Here, we build on this idea by leveraging large-scale motor unit recordings in humans to provide a probabilistic mapping of homonymous and heteronymous recurrent inhibition across six muscles and two contraction intensities during voluntary isometric contractions. Specifically, we implemented and validated a pipeline that integrates these experimental data with simulation-based inference (*20*, *21*). Using a simulation model that reproduces experimental motor unit behavior, we estimated the contributions of recurrent inhibition and higher-frequency components of common synaptic input to spiking activity. The model make all biological assumptions explicit by specifying the circuit components and their parameters, and the inference procedure returns a range of physiological parameter values compatible with the experimental data, along with their associated uncertainty. Our results revealed specific muscle-and intensity-dependent patterns of recurrent inhibition, challenging the assumption that recurrent inhibition changes uniformly and monotonically with contraction intensity across muscles.

## Results

### General framework

We simulated pools of 30 motor neurons using a leaky integrate-and-fire model, receiving recurrent inhibition via 10 Renshaw cells. Recurrent inhibition strength was set by a single parameter: the mean disynaptic inhibitory weight (motor neuron to motor neuron; Fig. 1B). Each motor neuron received three inputs: a low-frequency common input (0–5 Hz), independent noise (0–80 Hz), and a band-limited higher-frequency common input with tunable bandwidth and amplitude (Fig. 1A). Each simulation generated 100 s of spiking activity, representing an isometric submaximal contraction task (Fig. 1C). Full details are provided in the Methods section, and the code and parameter settings for all simulations are available on GitHub (https://github.com/FrancoisDernoncourt/Mapping_Recurrent_Inhibition).

**Figure 1.**
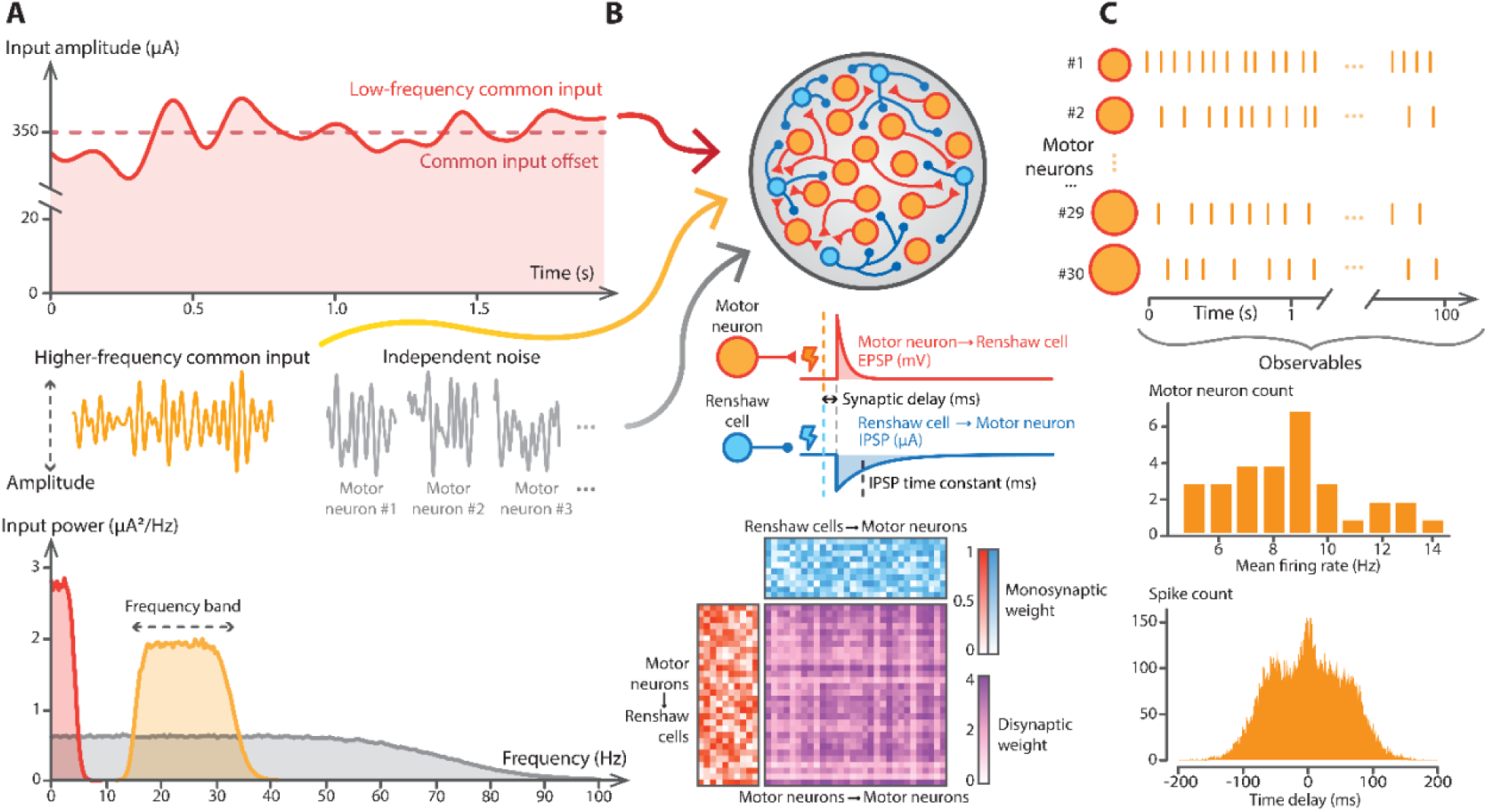
Simulation model overview. **(A)** Inputs were modelled as time-varying currents injected directly into leaky integrate-and-fire motor neurons. Within each simulated pool, all motor neurons received the same three components: a low-frequency common input (0–5 Hz), a band-limited higher-frequency common input, and a constant positive offset representing baseline excitation, i.e., common input offset. Each motor neuron also received independent noise (0–80 Hz). **(B)** Each motor neuron spike produced an excitatory postsynaptic potential (EPSP) of fixed amplitude and decay time constant in its target Renshaw cells, scaled by a monosynaptic weight. In turn, each Renshaw cell spike produced an inhibitory postsynaptic potential (IPSP) of fixed amplitude and adjustable time constant in its target motor neurons, also scaled by a monosynaptic weight. A single recurrent inhibition strength parameter set the mean disynaptic inhibitory weight between motor neurons. The corresponding monosynaptic weight matrices (motor neurons to Renshaw cells and Renshaw cells to motor neurons) were derived from this parameter (bottom panel; see Methods). **(C)** For the main analysis, we simulated 100 s of spiking activity in pools of 30 motor neurons with varying soma sizes and intrinsic properties. From the simulated spike trains, we extracted five observables, including mean firing rates and synchronization cross-histogram features. In this example, the cross-histogram represents a single comparison motor neuron, showing its conditional probability of spiking at a given time lag relative to spikes from all other reference motor neurons.

From the model outputs, we defined observable features from synchronization cross-histograms that served as proxies for recurrent inhibition and higher-frequency common input. To untangle the effects of recurrent inhibition from those of higher-frequency common input on these features, we used a simulation-based inference approach consisting of training a neural network on simulated data to probabilistically map these observable features to the latent physiological variables. Finally, we applied this trained network to experimental large-scale motor unit recordings to provide a probabilistic mapping of recurrent inhibition across six muscles, at two contraction intensities.

### Cross-histogram signatures of recurrent inhibition and higher-frequency common input

Synchronization cross-histograms were computed from simulated spike trains to represent the probability of a *comparison* motor neuron firing at specific time lags relative to a *reference* motor neuron. Specifically, for each *comparison*/*reference* pair, we measured the delays between each spike time of the *reference* motor neuron and the nearest preceding and following spikes of the *comparison* motor neuron. Counts were normalized to obtain a probability distribution. Finally, for each *comparison* motor neuron, we averaged its synchronization cross-histograms across all *reference* motor neurons to estimate the net inhibitory influence exerted by the other identified motor neurons of the pool.

A typical synchronization cross-histogram exhibits a curved, trapezoid-shaped baseline with a central peak flanked by troughs (Fig. 2). Each synchronization cross-histogram was fitted with a composite curve consisting of a *baseline* and a *peak-and-troughs* component (Methods). When applied to cross-histograms generated from simulated data, the goodness of fit was excellent (R^2^ = 0.97±0.04). We extracted three features from these fits. First, *trough area*, defined as the area where the histogram fell below the baseline. It quantifies the reduction in firing probability relative to baseline and served as a proxy for recurrent inhibition strength (Fig. 2A). Second, *trough timing,* defined as the lag corresponding to the minimum autocorrelation of the residuals relative to the baseline curve. This metric served as a proxy for the delay of recurrent inhibition, reflecting both the disynaptic delay and the time constant of the Renshaw-mediated inhibitory post-synaptic potential (IPSP; Fig 2A). Third, *peak height*, defined as the maximum positive amplitude of the *peak-and-troughs* component and used as an index of short-timescale synchrony, mainly arising from higher-frequency components of the common input (Fig. 2B).

**Figure 2.**
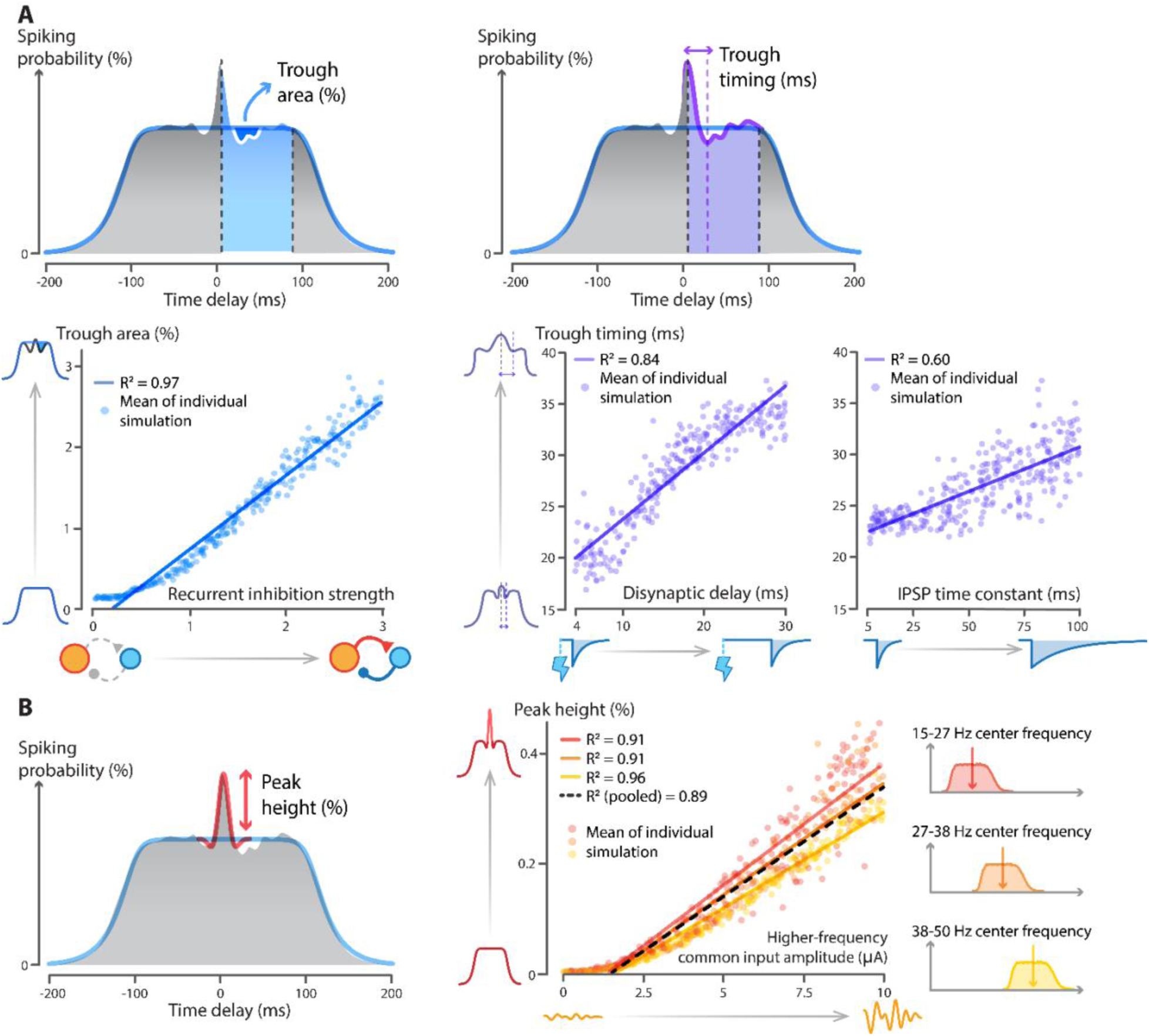
Synchronization cross-histogram features as proxies for recurrent inhibition and higher-frequency common input amplitude. **(A)** Example synchronization cross-histogram for one *comparison* motor neuron (participant #4, tibialis anterior at 10% maximal voluntary contraction (MVC) illustrating how features related to recurrent inhibition were defined. The *trough area* was calculated as the integrated reduction in spiking probability relative to the baseline curve (blue) following spikes of the *reference* motor neurons. The *trough timing* was defined as the lag at which the minimum autocorrelation of the cross-histogram residuals relative to the baseline curve occurred. The bottom panels show the relationships between these features and the corresponding ground-truth physiological parameters in the simulations. Each point represents the mean feature value across motor neurons from a single simulation, and the solid lines indicate linear regressions. **(B)** Example synchronization cross-histogram illustrating *peak height*, defined as the maximum positive amplitude of the *peak-and-trough* component of the fitted curve (red). The middle panel shows the relationship between *peak height* and the ground-truth amplitude of the higher-frequency common input across simulations. Each point represents the mean peak height across motor neurons from a single simulation. Colors denote three conditions with different center frequencies of the higher-frequency band. Linear regressions are shown for each condition, with the black dashed line indicating the regression across all conditions. IPSP: Inhibitory Postsynaptic Potential.

To validate that the three features tracked their intended physiological parameters, we simulated spike trains while independently varying recurrent inhibition and the higher-frequency component of the common input. We then quantified the linear relationships between each feature and its target physiological parameter. When varying recurrent inhibition strength across 300 simulations, linear regression of *trough area* against the ground truth explained 97% of the variance (R² = 0.97; Fig. 2A). At fixed inhibition strength, linear regression showed that *trough timing* explained 84% of the variance in ground-truth disynaptic delay (300 simulations; 4–30 ms delays; R² = 0.84; Fig 2A) and 60 % of the variance in the Renshaw-mediated IPSP time constant (300 simulations; 5-100ms; R² = 0.60; Fig 2A). For higher-frequency common input, we ran 1,000 simulations, jointly varying amplitude and frequency content of common input. The input was band-limited to a fixed 20-Hz bandwidth, with center frequencies spanning between 15 and 50 Hz (i.e., frequency bands from 5-25 Hz up to 40-60 Hz). Within each band, linear regression of *peak height* against input amplitude explained more than 91% of the variance (R² ≥ 0.91), and regressions remained high when pooling across bands (R²=0.89; Fig. 2B). Together, these results suggest that synchronization cross-histogram features can serve as proxies for estimating the amplitude of higher-frequency common input to motor neurons and the strength and delay of recurrent inhibition. However, our simulations also replicated prior observations that these proxies are not strictly mechanism-specific (Fig. 3); that is, the recurrent inhibition proxy is modulated by the frequency content of common input, and the proxy for high-frequency common input is, in turn, modulated by recurrent inhibition (*7*, *22*). This interaction makes it difficult to assign each feature exclusively to a single underlying mechanism, as discussed below.

**Figure 3.**
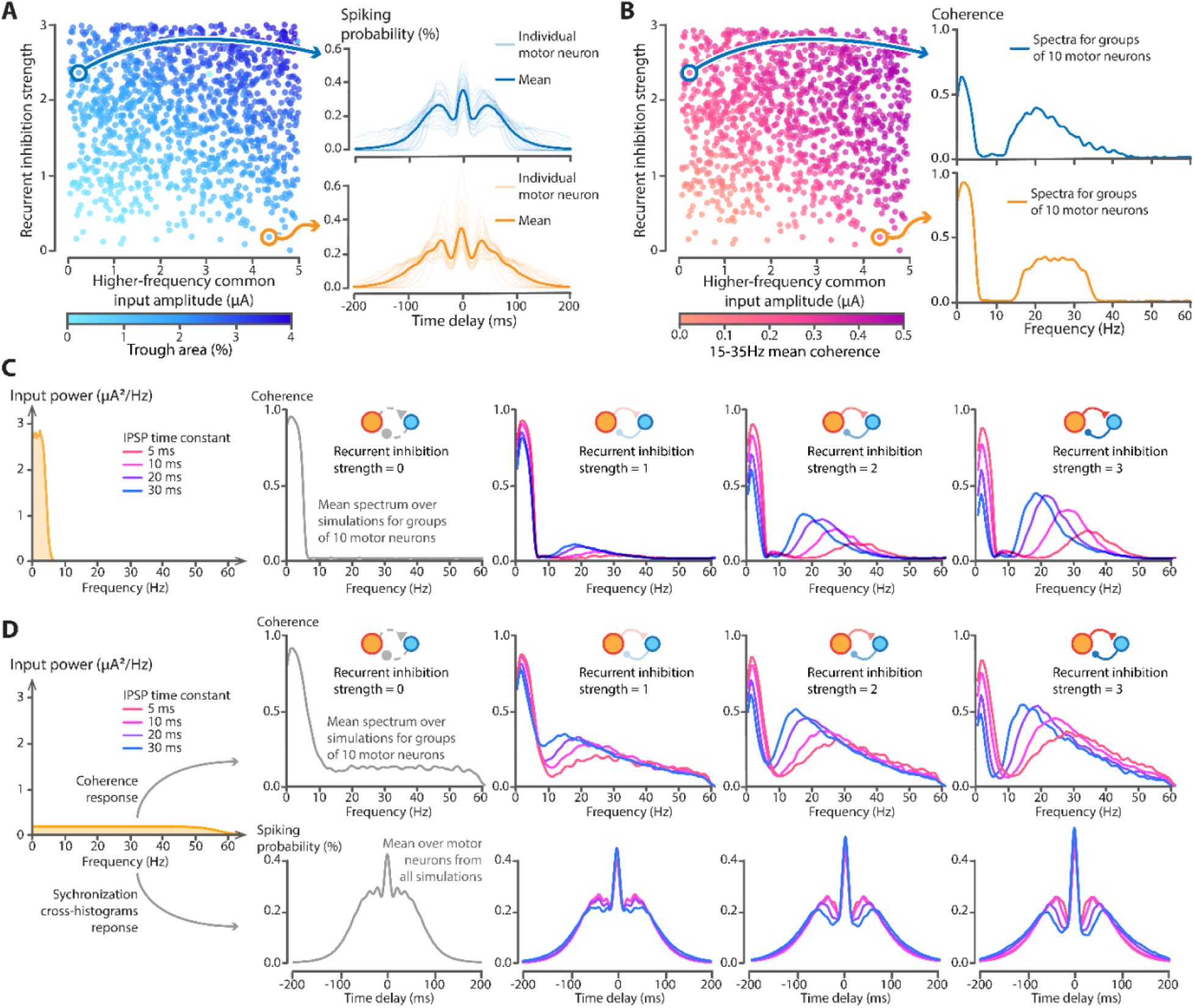
Interaction between common input and recurrent inhibition on synchronization cross-histogram features. **(A)** The scatterplot shows trough area for different combinations of higher-frequency common input amplitude (x-axis) and recurrent inhibition strength (y-axis). Each point corresponds to a simulation of 30 motor neurons. Synchronization cross-histograms are shown for two example simulations, where low-opacity lines represent each individual motor neuron and the high-opacity line represents their average. This example illustrates that two different combinations of higher-frequency common input amplitude and recurrent inhibition strength can produce similar synchronization cross-histograms. **(B)** Same layout as in panel A, with color coding representing the mean intramuscular coherence in the 15-35 Hz bandwidth between two groups of 10 motor neurons each. Intramuscular coherence spectra are shown for the same two example simulations as in panel A. **(C)** Effect of recurrent inhibition on coherence when only low-frequency common input and independent noise are delivered to the motor neuron pool. The left panel shows the power spectrum of the low-frequency common input (0–5 Hz) delivered to the pool, with the remaining four panels showing the mean intramuscular coherence between groups of 10 motor neurons each for different inhibitory post synaptic potential (IPSP) time constants (colors) and increasing recurrent inhibition strength (from left to right). **(D)** Effect of recurrent inhibition strength in the presence of broad-band common input. The left panel shows the power spectrum of the common input (0–50 Hz) delivered to the motor neuron pool, with total input power matching that in panel C. The top row shows mean intramuscular coherence between groups of 10 motor neurons for different IPSP time constants (colors) and increasing recurrent inhibition strength (left to right). The bottom row shows the corresponding synchronization cross-histograms for the same simulations.

### Recurrent inhibition as an intrinsic source of high-frequency synchrony

We used simulations to quantify cross-over effects between recurrent inhibition and the higher-frequency content of the common input on the proxy features, i.e., *trough area*, *trough timing,* and *peak height*, as well as on the intramuscular coherence spectrum of the cumulative spike trains. Using the in silico model introduced above, we generated a dataset of 3,000 simulations by varying recurrent inhibition strength and the amplitude of higher-frequency content of the common input (fixed 15-35 Hz band). To compare simulations at similar discharge rates, we varied common input offset (baseline excitation) and retained simulations with mean firing rate between 8 and 12 Hz. We observed that simulations with different input parameters could yield similar outputs. To illustrate, we selected two simulations with opposite parameters, i.e., low recurrent inhibition with large higher-frequency common input amplitude *vs* high recurrent inhibition with low higher-frequency common input amplitude. Despite these opposing parameters, the resulting synchronization cross-histograms (R²=0.79±0.20; Fig. 3A) and intramuscular coherence spectra were closely matched (R²=0.84±0.02; Fig. 3 B).

To explore this interaction further, we varied recurrent inhibition strength (three levels) and the Renshaw-mediated IPSP time constant (three levels) while supplying only low-frequency common input (0–5 Hz). Across these simulations, recurrent inhibition alone generated high-frequency synchrony, reflected as increased coherence at higher frequencies. Specifically, the IPSP time constant set the dominant frequency of the intramuscular coherence spectrum, whereas recurrent inhibition strength scaled the amplitude of the coherence (Fig. 3C). We then introduced a broadband common input (0–60 Hz at fixed amplitude) to examine the interaction between recurrent inhibition and common input. We ran nine scenarios: no recurrent inhibition, plus two inhibition strengths × 4 IPSP time constants. Recurrent inhibition behaved like a frequency-selective filter, i.e., it amplified oscillations near its intrinsic frequency, increasing intramuscular coherence at 15–35 Hz and the cross-histogram *peak height* (short-timescale synchrony), and suppressed slower fluctuations, reducing coherence below ∼15 Hz (Fig. 3D). In other words, recurrent inhibition amplifies input fluctuations that align with the recurrent loop’s intrinsic dynamics and suppresses slower fluctuations.

Overall, these simulations show that, even under simplified conditions, different combinations of recurrent inhibition and higher-frequency common input amplitude can generate similar synchronization cross-histograms and intramuscular coherence spectra. Disentangling the contribution of each mechanism from these features is therefore an ill-posed problem where multiple physiological parameter sets can account for the same observation. Such indeterminacy is common in neuroscience (*23*). To address this challenge, models can be used to identify the set of solutions compatible with the experimental data. Simulation-based inference provides a principled Bayesian framework for this purpose (*20*) and has been successfully applied in neuroscience (*21*, *24*, *25*). In the current study, we implemented and validated a simulation-based inference workflow to infer plausible distributions of recurrent inhibition strength and higher-frequency common input amplitude from experimentally observable features, as detailed in the section below.

### Simulation-based inference to quantify recurrent inhibition

To dissociate the effects of recurrent inhibition from those of higher-frequency common inputs on synchronization cross-histogram features, we used simulation-based inference with our in silico model as the simulator. First, we defined ranges for five physiological parameters: (1) baseline excitation (common input offset), (2) recurrent inhibition strength, (3) amplitude of the higher-frequency component of the common input, along with its frequency band described by (4) its central frequency and (5) its half-width. These ranges defined the *priors* and were chosen so that simulated observations covered the range of experimental observations (Methods; Fig. S1). All other parameters were held constant across simulations. Second, we sampled sets of parameters from these *priors* and generated simulated spike trains (12,000 simulations for single motor neuron pools and 20,000 simulations for pairs of synergistic motor neuron pools – see Methods). From each simulation, we computed four observables: the three cross-histogram features (*trough area*, *trough timing*, *peak height*) and the mean firing rate. This yielded a large training dataset consisting of pairs of parameter sets and their corresponding observed features. Third, we trained a neural density estimator (a neural network that outputs probability distributions) on this dataset to learn a probabilistic map between parameters and observed features. This mapping can then be inverted to infer which physiological parameter values are compatible with a given set of observed features. Finally, we fed summary statistics (mean, median, standard deviation, interquartile range) of experimentally observed features into the trained neural density estimator, yielding a *posterior* distribution over physiological parameter combinations (see the next section for the experimental results). In other words, the estimator returned combinations of parameters (baseline excitation, recurrent inhibition strength, higher-frequency common input amplitude and frequency) that can explain the experimentally observed features (*trough area*, *trough timing*, *peak height*, mean firing rate), with quantified uncertainty (Fig 4A; Methods). As recommended in the literature, we validated this pipeline on simulated data with known ground truth and on experimental data (*25*, *26*).

**Figure 4.**
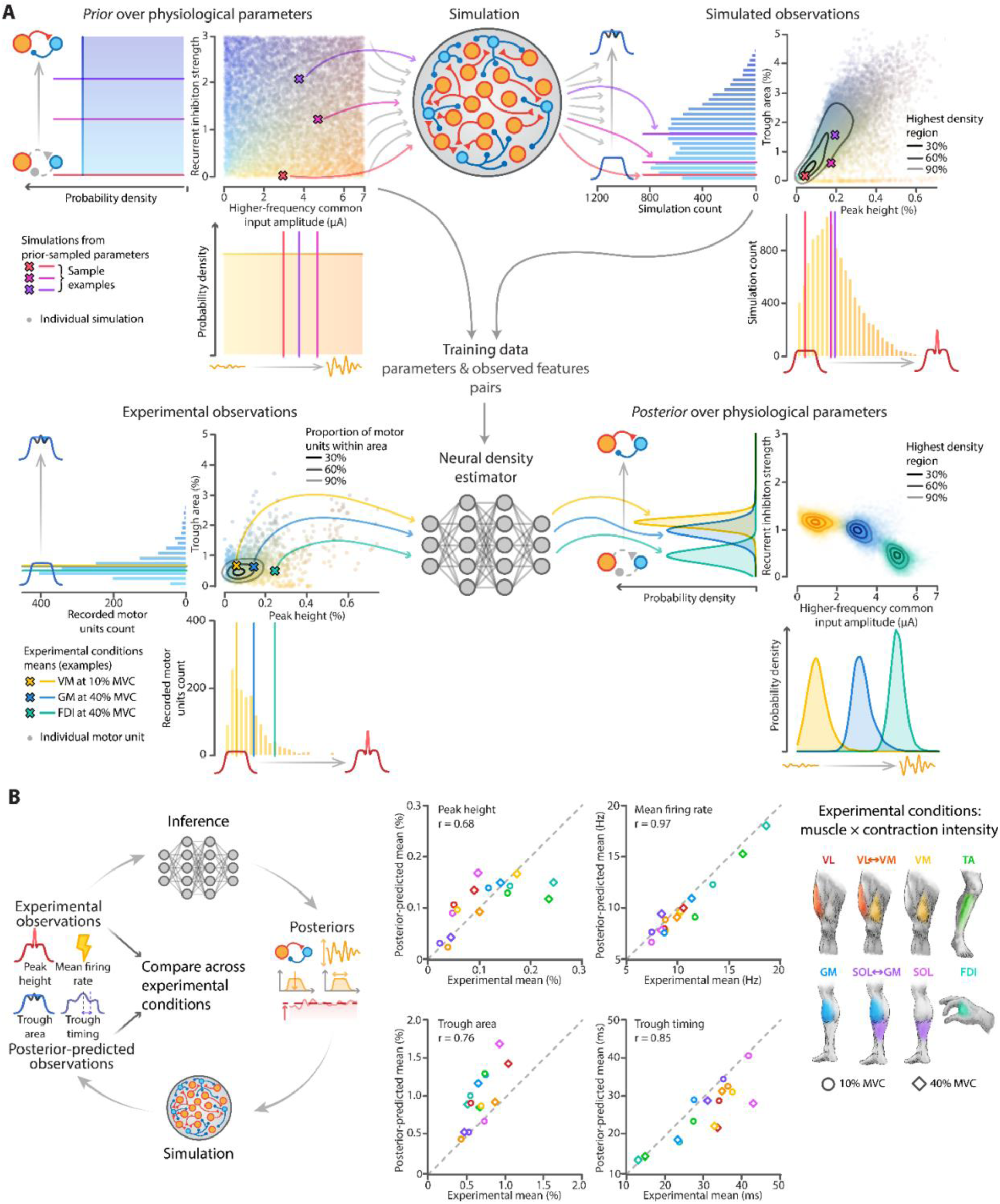
Simulation-based inference pipeline and validation. **(A)** Physiological parameters were sampled from their *prior* distributions. The scatterplot (top left) shows the 12,000 *prior* draws used for simulations. For visualization, only two parameters are shown (recurrent inhibition strength and higher-frequency common input amplitude), with marginal density plots showing their individual *prior* distributions. Each parameter set was used in the simulation model to generate motor neuron spike trains, from which observable features were extracted. Only *trough area* and *peak height* are illustrated (top right panels). Three example parameter sets are shown in different colors. The resulting parameter–feature pairs were used to train a neural density estimator to learn the inverse mapping from features to parameters. Given a set of features derived from experimental data, the neural density estimator returns a *posterior* distribution over the physiological parameters (bottom panels). The 2D density plot (bottom right) shows the joint *posterior* over recurrent inhibition strength and higher-frequency common input amplitude for three example conditions, with marginal density plots showing the corresponding *posteriors* for each parameter. **(B)** *Posterior* predictive checks on experimental data. The schematic (left) summarizes the procedure. For each experimental condition (muscle × contraction intensity), the experimentally observed features were fed into the trained neural density estimator, which returned the corresponding *posteriors* over physiological parameters. These inferred parameters were then used in simulations to assess how well they reproduced the experimentally observed features. Each of the four middle panels shows the relationship between the experimental mean of a feature and its corresponding *posterior*-predicted mean across all conditions (muscle × contraction intensity). VL: Vastus lateralis; VM: Vastus Medialis; GM: Gastrocnemius Medialis; SOL: Soleus; FDI: First Dorsal Interosseous, TA: Tibialis anterior; MVC: Maximal voluntary contraction.

We first evaluated the density estimator on simulated data held out from training (10% of the dataset). This provided an upper bound on performance under correct model specification, as the held-out data were generated by the same model as used to train the neural density estimator (Supplementary Fig. S2). Specifically, for each of the five [0,1]-normalized physiological parameters, we quantified the estimator’s *accuracy* (how well it recovers the ground truth) and its *calibration* (whether its *posteriors* are over-or under-confident). For accuracy, we computed the root mean square error (RMSE) and Pearson’s correlation coefficient between a parameter’s ground truth values and its *posterior* modes (highest-probability values according to the *posterior*) across all simulations (Supplementary Fig. S2A). When considering baseline excitation, recurrent inhibition strength, and higher-frequency common input amplitude, this yielded low error levels (RMSE ≤ 0.124 compared with RMSE ≅ 0.41 expected from *prior*; r ≥ 0.90 compared with r = 0 expected from *prior*). However, when considering the frequency band center and half-width, the accuracy was low (RMSE ≥ 0.33; r ≈ 0). Calibration was assessed using a coverage curve. We incrementally increased the nominal highest-*posterior*-density mass from 0 to 100% and, at each step, measured the proportion of held-out ground-truth values that fell within the corresponding highest-*posterior*-density interval (Methods). A perfect calibration would follow the identity line. Our curves were slightly below this line, indicating mild overconfidence, but overall calibration remained strong (mean calibration error ≤ 3.9% for all parameters; supplementary Fig. S2B). Together, these analyses performed on simulated data demonstrate that while detailed frequency content could not be recovered accurately using our approach, the parameters most relevant to our research question, including recurrent inhibition strength, are estimated accurately with well-quantified uncertainty. Of note, supplementary Fig. S3 illustrates how simulation-based inference resolves the indeterminacy example shown in Fig. 3A–B, and how the degree of uncertainty of the *posterior* depends on the information content of the observed features.

We further tested the accuracy of our approach with *posterior* predictive checks on the experimental dataset (six male participants, six muscles, and two intensities). For each muscle × intensity combination (hereafter “condition”), we used the neural density estimator trained on simulated data to infer a *posterior* distribution over the physiological parameters, i.e., baseline excitation, recurrent inhibition strength, higher-frequency common input amplitude, central frequency and frequency band half-width. Then, we sampled 100 parameter sets from this *posterior* distribution, and we used those parameter sets to simulate motor neuron spike trains, leading to 1600 *posterior*-predictive simulations ([6 muscles + 2 synergist pairs] × 2 intensities × 100). From these spike trains, we constructed synchronization cross-histograms, extracted three features (*trough area*, *peak height, trough timing*), added the mean firing rate as a fourth feature, and computed the mean of each feature across motor neurons for every simulation. This yielded a distribution of *posterior*-predicted feature means for each condition. We standardized each feature to the experimental data so that an RMSE of 1 corresponds to an average prediction error equal to one standard deviation of the experimental values for that feature. We then compared, for each condition, the mean of the *posterior*-predicted distribution of feature means with the experimentally observed feature mean (Fig. 4B). Accuracy was very good for mean firing rate (RMSE=0.32), moderate for *peak height* (RMSE=0.73) and *trough timing* (RMSE=0.76), and poor for *trough area* (RMSE=1.41). However, errors were mostly due to systematic bias of the *posterior* predictions (overestimation of *trough area* and underestimation of *trough timing*), whereas the relative difference across conditions was well preserved. This is reflected in the high correlation coefficients between experimental and *posterior*-predicted feature means across conditions (mean firing rate: r=0.97, *trough area*: r=0.76; *peak height*: r=0.68; *trough timing*: r=0.85), indicating that the estimator reliably captures between-condition differences even when absolute levels are biased.

In summary, these validation checks support the validity of the simulation-based inference pipeline: it recovered key parameters on held-out simulations with good calibration, and *posterior* predictions qualitatively reproduced the patterns of experimental features across conditions. We therefore used this approach to map homonymous and heteronymous recurrent inhibition in human spinal motor neurons innervating different muscles.

### Experimental data

The experimental setup and main processing steps are illustrated in Fig. 5. After excluding 16 motor units (<1% of the total) for poor synchronization cross-histogram fits or low spike counts (Methods), we retained 1669 unique motor units over the six participants, with an average of 109±76 motor units per muscle and contraction intensity (range: 27 for VM at 40% MVC to 254 for GM at 40% MVC). The full dataset (raw signals and motor unit spike trains) is available at: https://figshare.com/s/c4da3eb57c88a9b07ddc.

**Figure 5.**
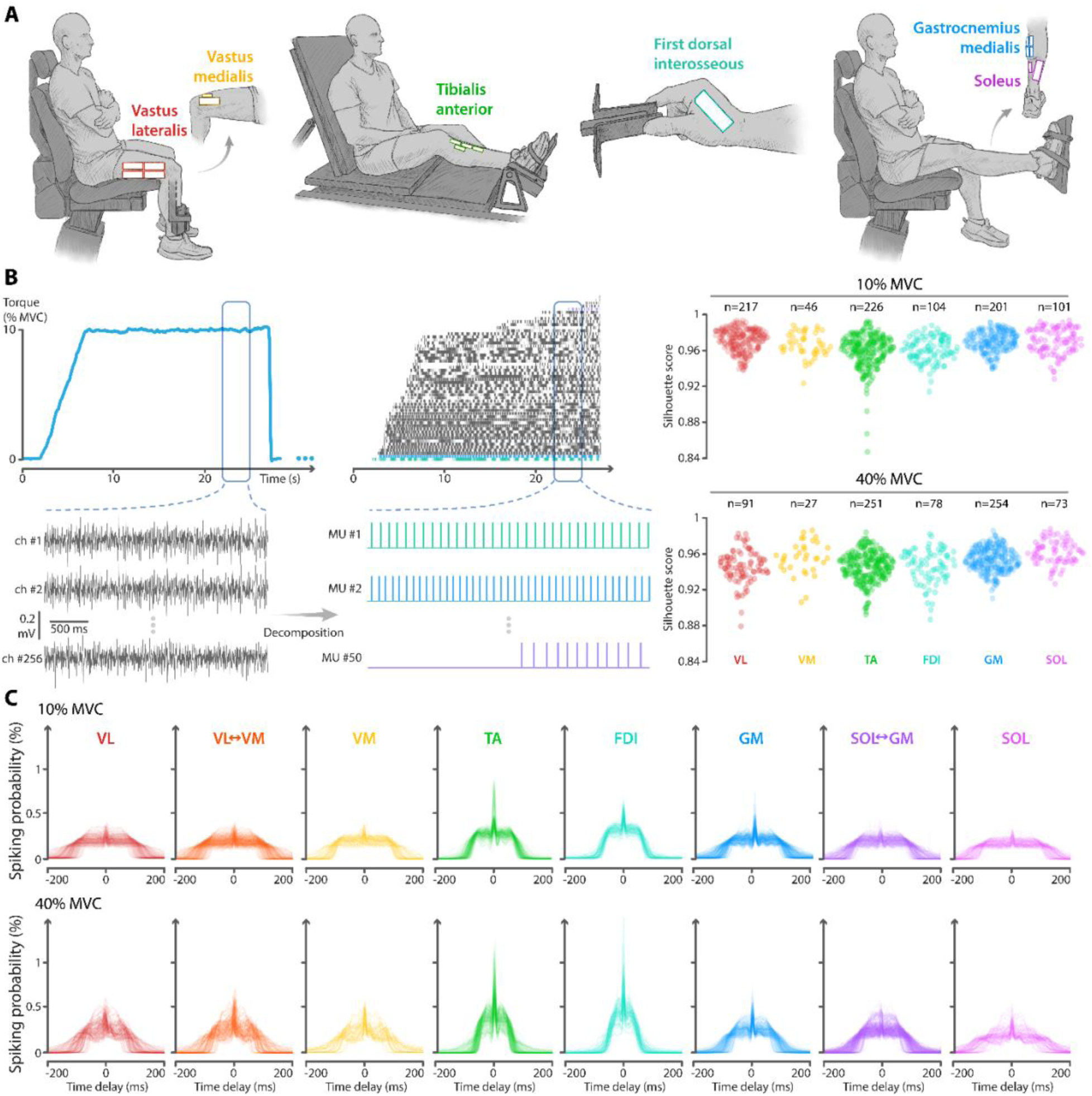
Experimental setup and analysis. **(A)** Experimental setup for the four tasks (from left to right): knee extension, dorsiflexion, pinch, and plantarflexion. **(B)** Example of the motor unit decomposition pipeline, illustrated for the knee-extension task at 10% MVC in participant #1. Insets show the produced torque, an example of interference electromyographic signals, and the identified spike trains from three motor units. The right panels show the distribution of median silhouette scores for each identified motor unit across all participants after decomposition and manual editing. Silhouette scores were computed over non-overlapping 1-s windows. They quantify the reliability of motor unit spike identification. *n* indicates the number of retained motor units for each muscle and contraction intensity. **(C)** Full set of experimental synchronization cross-histograms, grouped by condition (muscle × contraction intensity) and pooled across participants. Each thin line represents one motor unit’s synchronization cross-histogram. MU: motor unit; VL: Vastus lateralis; VM: Vastus medialis; GM: Gastrocnemius medialis; SOL: Soleus; FDI: First dorsal interosseous, TA: Tibialis anterior; MVC: Maximal voluntary contraction.

Using spike trains recorded over 200 s per participant, we constructed synchronization cross-histograms for each motor unit, fitted them with the composite curve (R²=0.99±0.02), and extracted the features used for inference, i.e., mean firing rate, *trough area*, *trough timing*, and *peak height*. The simulation-based inferences used summary statistics of these feature distributions, aggregated across all motor units from all participants for each condition (muscle × intensity). Note that the approach is readily applicable at the single-participant level. Here, we pooled data across participants to obtain more generalizable comparative inferences. We focused on two physiological parameters: recurrent inhibition strength and higher-frequency common input amplitude. For each condition, Fig. 6A shows the *posterior* distribution of each physiological parameter (marginal *posteriors*) as well as their joint *posterior* distribution. We report between-condition contrasts as pΔ, defined as the *posterior* probability that the specified inequality (e.g., A>B) is true (Fig. 6B). Thus, “A>B: pΔ=0.9” indicates a 90% probability that A exceeds B. When reporting heteronymous recurrent inhibition between synergist muscles, we use a single bidirectional notation (VL↔VM and GM↔SOL), because the directional *posteriors* (VL→VM *vs* VM→VL; GM→SOL *vs* SOL→GM) were nearly identical (Methods). Of note, we also performed an additional analysis in which motor unit spike trains were independently shuffled, thereby preserving firing rate statistics while disrupting synchronization cross-histogram features (Fig. S4). This analysis was used to assess whether the *posterior* distributions differed from what would be expected by chance under this shuffling procedure.

**Figure 6.**
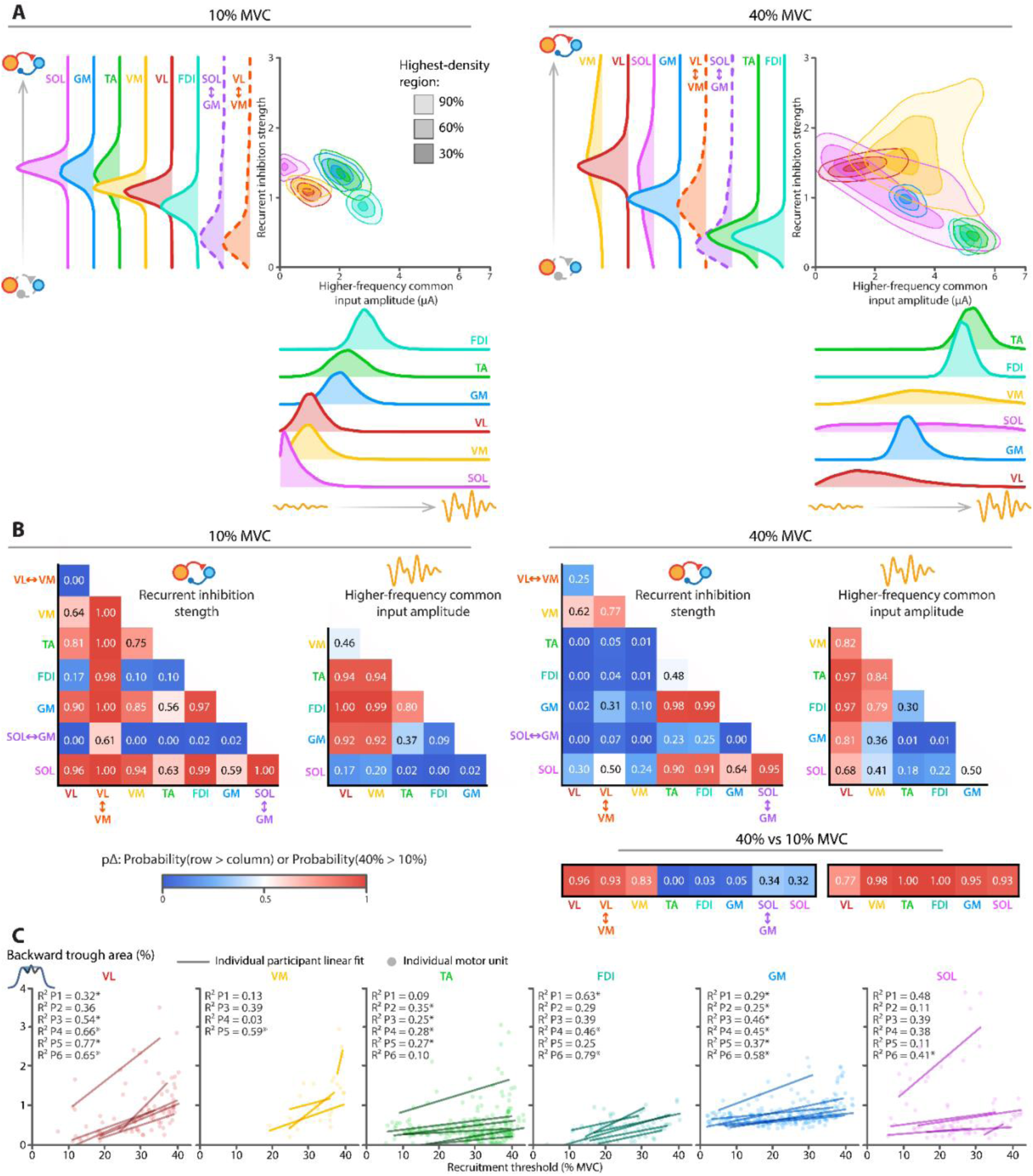
Results from simulation-based inference. **(A)** *Posterior* distributions of recurrent inhibition strength and higher-frequency common input amplitude for each muscle or muscle pair at 10% (left) and 40% (right) of maximal voluntary contraction (MVC). The 2D density plots show the joint *posteriors* over these two parameters, and the 1D density plots show the marginal *posteriors*. Inference was performed for each muscle × contraction intensity condition using the summary statistics derived from the four observed features. Note that higher-frequency common input amplitude was inferred only for individual muscles, which explains the absence of between-muscle conditions. **(B)** Pairwise comparisons of the inferred *posteriors* for recurrent inhibition strength and higher-frequency common input amplitude. In each matrix, the color scale represents the probability that the marginal *posterior* for the row muscle exceeds that for the column muscle within a given intensity (10% on the left, 40% on the right), or that the *posterior* at 40% MVC exceeds that at 10% MVC for a given muscle (bottom right). **(C)** Scatterplots show the backward trough area (a proxy for delivered recurrent inhibition, as explained in the main text) as a function of recruitment threshold for contractions performed at 40% MVC. Each point represents an individual motor neuron, and lines represent regressions fitted separately for each participant. Asterisks mark regressions that were statistically significant (p<0.05, Benjamini–Hochberg correction). VL: Vastus lateralis; VM: Vastus medialis; GM: Gastrocnemius medialis; SOL: Soleus; FDI: First dorsal interosseous; TA: Tibialis anterior; MVC: Maximal voluntary contraction.

We first report results obtained at 10% MVC. Consistent with previous observations that recurrent inhibition is low or absent in intrinsic hand muscles (*6*, *27*), homonymous recurrent inhibition was lowest in FDI, with a high probability of being lower than in the calf muscles (all pΔ>0.97). In contrast, lower leg muscles (GM, TA, SOL) exhibited the highest recurrent inhibition overall, with values likely higher than in thigh muscles (VL and VM; all pΔ≥0.75). Finally, heteronymous recurrent inhibition (i.e., between synergist muscles) was negligible at this low contraction intensity, as the *posterior* distributions overlapped with those obtained from the shuffled dataset (Fig. S4A).

When contraction intensity increased from 10% MVC to 40% MVC, recurrent inhibition changed in a muscle-specific manner. In line with reports that recurrent inhibition decreases with increasing force (*16*, *27*, *28*), homonymous recurrent inhibition decreased in GM, TA, and FDI (all pΔ≥0.95). In contrast, the vastii exhibited the opposite pattern, with a high probability of increase in both homonymous and heteronymous recurrent inhibition (all pΔ≥0.83; Fig. 6B). Changes in SOL were inconclusive. As a result, at 40% MVC, recurrent inhibition was higher in thigh muscles (VL and VM) than in GM, TA, and FDI. While GM↔SOL heteronymous recurrent inhibition remained negligible, VL↔VM heteronymous recurrent inhibition became substantial at 40% MVC, with a high probability of exceeding homonymous inhibition in FDI (pΔ=0.89).

The amplitude of the higher-frequency common input was generally greater in TA and FDI than in the other muscles at both 10% and 40% MVC. At 10% MVC, TA and FDI values likely exceeded those of SOL: VM, and VL (all pΔ ≥ 0.94); and at 40% MVC, they likely exceeded those of VL and GM (all pΔ ≥ 0.97). Across muscles, there was a high probability of an increase in the amplitude of higher-frequency common input from 10% MVC to 40% MVC.

Together, our results replicate well-established findings while adding key insight. Specifically, we show muscle-specific changes in recurrent inhibition with increasing contraction intensity, leading to marked differences across muscles at 40% MVC: recurrent inhibition was overall greater in the vastii, and VL↔VM heteronymous inhibition exceeded homonymous recurrent inhibition in FDI and TA.

Finally, we conducted additional analyses to assess whether recurrent inhibition is homogeneously distributed within the motor neuron pool. Prior work suggests that recurrent inhibition depends on motor neuron size (*22*, *29–31*), but direct confirmation in humans is lacking. Here, we leveraged our large-scale motor unit recordings to examine size dependence in recurrent inhibition using a per-neuron proxy feature for *delivered* inhibition: the *trough area* computed at negative lags (*backward* in time) rather than positive lags. We made this choice to minimize confounding effects arising from intrinsic motor neuron properties. Indeed, lower-threshold motor neurons, having higher membrane resistance, are more sensitive to inhibitory inputs. Consequently, the *trough area* computed forward in time, as in the other analyses, can be larger in these neurons than in higher-threshold neurons, not because they receive greater inhibition, but because of their intrinsic properties. In contrast, the *trough area* assessed at negative lags reflects inhibition that a *comparison* motor neuron *delivers* to all other neurons. Because synchronization cross-histograms are averaged across many motor neurons, intrinsic property-dependent effects are largely cancelled out, allowing us to approximate the level of recurrent inhibition delivered by each *comparison* motor neuron as a function of its size. We report this analysis for the contractions performed at 40% of MVC, where a broader range of motor neuron sizes is recruited. Within each muscle and participant, recruitment threshold (a proxy for motor neuron size) was positively related to *backward trough area* at 40% MVC (R²=0.38±0.19), with 20 out of 34 linear regressions reaching statistical significance (Fig. 6C). This indicates that higher-threshold (larger) neurons tend to induce more recurrent inhibition than lower-threshold (smaller) neurons.

## Discussion

Although Renshaw cell–mediated spinal recurrent inhibition was among the first spinal circuits to be described (*4*, *5*), its functional role and regulation remain poorly understood. In humans, direct measurements are not feasible, and recurrent inhibition can only be inferred indirectly, with inherent limitations. In this study, we developed and validated a framework that combines experimental data with simulation-based inference to estimate homonymous and heteronymous recurrent inhibition from motor unit spiking behavior across six muscles during isometric contractions at two force levels. While several of our findings are consistent with previous observations in humans and animal models, others reveal a more nuanced pattern of variation in recurrent inhibition across muscle groups and contraction intensities than is typically assumed.

We estimated recurrent inhibition from synchronization cross-histograms constructed from motor neuron spike trains. However, simulations revealed that these cross-histograms are also influenced by high-frequency components of the common synaptic input. To disentangle these effects, we developed a simulation-based inference framework (*20*, *21*), in which a neural density estimator was trained on simulated data. Feeding the experimentally observed features into this model returned probability distributions over recurrent inhibition strength. In other words, this approach allows us to interpret the experimental data in a principled and testable manner, with all assumptions made explicit. Unlike stimulation paradigms (*16*, *17*, *28*), our approach estimates recurrent inhibition from naturalistic behavior, i.e. voluntary contractions, and can be applied to any muscle from which motor unit spike trains can be identified.

To interpret our findings appropriately, several methodological considerations must be addressed. First, a key limitation of our approach lies in the simplified nature of the simulation model. Based on leaky integrate-and-fire neurons, the model reproduces realistic motor neuron activity while capturing the effects of recurrent inhibition and common input. However, it deliberately excluded other spinal pathways and state-dependent nonlinearities, such as those arising from inward currents. While these simplifications are reasonable for short, non-fatiguing submaximal isometric contractions at constant force, where proprioceptive feedback (*32*) and history-dependent motor neuron behavior (*33*, *34*) are expected to remain approximately stable, they limit the generalizability of the results. In particular, the findings do not directly extend to dynamic contractions, in which the omitted mechanisms are likely to play a larger role. A similar limitation applies to higher contraction intensities. Because motor unit decomposition deteriorates at higher activation levels, leading to lower motor unit yields, we limited the intensity to 40% MVC. Accordingly, our conclusions should not be extrapolated beyond this range. Second, all inferences should be interpreted in the context of the model’s level of abstraction. Specifically, because the model is an abstract approximation of the underlying biology, inferred recurrent inhibition represents a model-defined effective quantity. That is, it indicates that the observed patterns of motor unit synchronization are compatible with Renshaw-mediated inhibition, rather than providing a direct measure of a physiological mechanism. Third, the reliability of synchronization cross-histogram features depends on spike count: lower motor neuron yields and shorter recordings produce noisier histograms. Importantly, a sensitivity analysis showed that lower spike counts reduce the likelihood that a recording meets inclusion criteria but does not substantially affect the accuracy of the *posterior* estimates once quality-control thresholds are met (Methods; Fig. S5). Because we analyzed long contractions (a total of 200 s) and used multiple electrode grids (up to 256 electrodes), enabling the identification of a relatively large number of motor neurons (e.g., a mean of 42 per participant in GM at 40% MVC), our experimental data exceed the minimum requirements for reliable inference (Fig. S5). Fourth, we acknowledge that experimental data were obtained exclusively from male participants. This was primarily due to technical limitations, as we were unable to reliably decode a sufficient number of motor units from the vastii and soleus muscles in female participants (pilot data) to support the present analyses. Finally, our study does not provide direct biological validation against experimental ground truth. Because the inference relies on cross-histograms, our method requires simultaneous recordings from a relatively large number of motor neurons. Therefore, direct biological validation would require datasets combining (i) simultaneous recordings of a large number of motor neurons with either (ii.a) direct measurements of Renshaws cell output or (ii.b) selective control of recurrent inhibition. To our knowledge, no existing datasets provide both components. Despite these limitations, the validation of the generative model, together with our ability to reproduce well-established features of recurrent inhibition — including several observed directly in animal models — give us confidence that the aforementioned limitations do not substantially affect the interpretation of our results.

Our results reveal a low level of recurrent inhibition in FDI, consistent with cross-species evidence that intrinsic hand muscles display little to no recurrent inhibition compared with proximal muscles (cat (*35*, *36*); human (*13*, *27*); primate (*6*)). Second, we observed a decrease in recurrent inhibition with contraction intensity in some lower-leg muscles, consistent with classic findings in humans (*16*, *27*, *28*). Direct recordings in reduced animal preparations also indicate frequency-dependent short-term depression in Renshaw-cell responses during prolonged stimulation trains, which may diminish their inhibitory influence (*37*). Third, the *trough timing* we recovered (e.g., 33.6±13.3 ms for VL at 40% MVC) falls within the range reported previously (∼20–40 ms (*17*, *22*, *38*)). Fourth, size-dependent trends were observed in our data, consistent with direct measurements in animal (*29–31*) and indirect observations in humans (*22*), indicating that larger motor neurons deliver stronger recurrent inhibition than smaller ones.

Beyond replicating classical results, our approach revealed new results that challenge long-held assumptions. Specifically, unlike the classical pattern observed in lower-leg muscles, both homonymous and heteronymous recurrent inhibition increased in the vastii with contraction intensity. Of note, because Renshaw cells activity is driven by motor neuron axon collaterals, this increase in recurrent inhibition with force is the logical consequence of increased motor neuron firing rate. Conversely, the decrease in recurrent inhibition classically observed in other muscles, and confirmed in our data, likely reflects a supraspinal downregulation of Renshaw cells (*39*). Taken together, these results reveal fundamental differences in the neural control of distinct motor neuron pools: in some muscles (e.g., GM, TA, Fig. 6A-B), inhibition is reduced as excitatory drive increases (a “push-pull” scheme (*40*)), whereas in others (e.g. VL, VM; Fig. 6A-B), inhibition is maintained or even increased in parallel with excitation (a “balanced excitation-inhibition” scheme (*41*)). These control schemes, mediated in part by recurrent inhibition, may help explain the well-documented differences in motor unit behavior across motor pools and contraction intensities. For example, in some muscles such as the FDI, most motor units are recruited early during a ramp-up contraction, and force is primarily modulated through rate coding (*1*, *42*). In contrast, in the VL muscle, force modulation is primarily achieved through motor unit recruitment, with only limited modulation of firing rate (*1*, *42*, *43*). This latter pattern is consistent with a balanced excitation-inhibition regime, in which increased excitation is accompanied by a parallel increase in inhibition (*44–46*). Of note, if later recruited, larger neurons also exert stronger inhibitory influence on earlier-recruited smaller neurons, this rate-limiting effect should become even more pronounced. Our finding that recurrent inhibition in the VL and VM is stronger at 40% MVC than at 10% MVC supports the idea that these muscles operate closer to a balanced excitation-inhibition regime than the other muscles examined. Between-muscle differences in recurrent inhibition patterns may also help explain variations in motor unit hysteresis and input-output nonlinearities. Because inhibition limits the amplification effect of persistent inward currents (*47–50*), the balanced excitation-inhibition pattern in vastii predicts smaller hysteresis and a flatter frequency-current curve compared with other muscles (*19*, *51*), particularly at higher contraction intensities, as confirmed by recent findings (*34*, *52*). In summary, our estimates of recurrent inhibition offer a plausible mechanism for the reported differences in motor unit behavior across muscles and raise questions about the functional relevance of the distinct inhibition scheme observed in the vastii.

The knee extensors have a role in joint-protection that goes beyond generating knee extension torque. In particular, coordinated VL and VM activity ensures mediolateral tracking of the patella (*53*, *54*), while quadriceps-hamstring co-contraction limits anterior tibial translation and reduces cruciate ligament loading (*55*, *56*). We propose that these objectives are at least partly implemented by fast, low-level spinal circuits in which recurrent inhibition plays a key role. In addition to reducing agonist (homonymous) motor neuron activity, recurrent inhibition also increases antagonist activity by inhibiting Ia inhibitory interneurons (*57*, *58*), which has led to the suggestion that recurrent inhibition may contribute to the control of joint stiffness (*59*). This interpretation fits within the framework that considers spinal circuitry as a predictive–comparator system (*8*, *10*). Specifically, these latter studies suggest that alpha motor neurons act as comparators, integrating excitatory proprioceptive (“instructive”) Ia input with a negative efference copy (the “predictive” inhibitory input) delivered by Renshaw cells to compute a local sensory prediction error. If this error term is defined over the full set of task objectives, including minimizing internal joint stress for example, recurrent inhibition may become an adaptive, rapid error-correction mechanism that can tune agonist (by inhibition) and antagonist (by disinhibition) motor neuron activity to regulate internal joint stress. Consistent with this idea, Renshaw cells are required for the behavioral expression of a learned association between a specific limb position and a noxious stimulus (*9*), where the relevant error signal can be viewed as the limb approaching a configuration associated with nociceptive input. Overall, the increase in homonymous and heteronymous recurrent inhibition we observed in the vastii at higher force levels may reflect a greater contribution of joint-protective control mechanisms implemented at the spinal level. This interpretation is further supported by reports of stronger recurrent inhibition during eccentric than during concentric or isometric contractions (*60*), conditions characterized by higher internal joint stress.

Beyond the physiological insights, this work introduces a framework for inferring recurrent inhibition from motor unit activity during voluntary contractions, which was not achievable with existing approaches. The full pipeline, including the spiking network simulator, feature-extraction scripts, and trained neural density estimators, is openly available. Researchers can apply the current model to compatible datasets, provided that the observed features fall within the range covered by the simulated training data. For new contexts (e.g., different muscles, tasks, or clinical populations) or to investigate other spinal circuits, the framework can be adapted by modifying the simulation model and/or *priors*, generating a new simulation dataset, and retraining the neural density estimator.

## Materials and Methods

### Simulation model

We simulated the spiking activity of motor neurons with varying recurrent inhibition and different characteristics of common synaptic inputs. To this end, we used a simulator for spiking neural networks (Brian 2 (*61*)) to implement our models in Python. The following section presents the main features of the simulation framework. The complete simulation code and full list of parameters are publicly available at: https://github.com/FrancoisDernoncourt/Mapping_Recurrent_Inhibition

#### Motor neuron model

We modeled each motor neuron as a leaky integrate-and-fire neuron with a resting membrane potential and reset voltage 𝑉_𝑟𝑒𝑠𝑡_ = 0 mV, and a firing threshold 𝑉_𝑡ℎ𝑟𝑒𝑠ℎ_ = 10 mV. The membrane potential change at each time step, 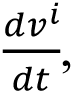 was computed using Euler integration (time step 𝑑𝑡 = 0.1ms) according to the equation below:

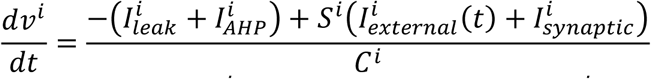

Where 𝑖 denotes motor neuron index, 𝐶^𝑖^ is membrane capacitance, 𝑆^𝑖^ is an input scaling factor, 𝐼^𝑖^𝑙𝑒𝑎𝑘 is the motor neuron leak current, 𝐼^𝑖^𝑒𝑥𝑡𝑒𝑟𝑛𝑎𝑙 (𝑡) is the net external input current (baseline input + common input + independent input), and 𝐼^𝑖^is the Renshaw-mediated inhibitory synaptic current. After a spike, 𝑣^𝑖^ was reset to 𝑉_𝑟𝑒𝑠𝑡_, an absolute 5 ms refractory period was enforced (𝑣^𝑖^ clamped at 𝑉_𝑟𝑒𝑠𝑡_), and a decaying afterhyperpolarization 𝐼^𝑖^_*AHP*_ current was induced. Each current was calculated as follows:

- 𝐼^𝑖^𝑙𝑒𝑎𝑘 = 𝑔^𝑖^𝑙𝑒𝑎𝑘 (𝑣^𝑖^ − 𝑉𝑟𝑒𝑠𝑡) with 𝑔^𝑖^𝑙𝑒𝑎𝑘 being the leak conductance, calculated as 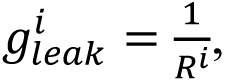 where 𝑅^𝑖^ is the membrane resistance.
- 𝐼^𝑖^𝐴𝐻𝑃 = 𝑔^𝑖^𝐴𝐻𝑃 (𝑣^𝑖^ − 𝑉_𝐴𝐻𝑃_) where 𝑔^𝑖^𝐴𝐻𝑃 is the afterhyperpolarization conductance. This conductance increased by a fixed increment Δ𝑔_𝐴𝐻𝑃_ (1 mS) at each spike and decayed over time according to 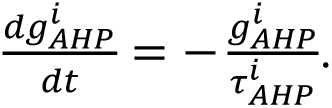 was set to-13.3 mV, corresponding to the-90 mV potassium reversal potential under our convention 𝑉_𝑟𝑒𝑠𝑡_ = 0 mV and 𝑉_𝑡ℎ𝑟𝑒𝑠ℎ_ = 10 mV.
- The net external input current was defined as 𝐼^𝑖^𝑒𝑥𝑡𝑒𝑟𝑛𝑎𝑙 (𝑡) = 𝑚𝑎𝑥(𝐼_𝑏𝑎𝑠𝑒𝑙𝑖𝑛𝑒_ + 𝐼_𝑐𝑜𝑚𝑚𝑜𝑛_(𝑡) + 𝐼^𝑖^𝑖𝑛𝑑𝑒𝑝𝑒𝑛𝑑𝑒𝑛𝑡 (𝑡) − 𝐼^𝑖^𝑟ℎ𝑒𝑜𝑏𝑎𝑠𝑒, 0), where 𝐼_𝑏𝑎𝑠𝑒𝑙𝑖𝑛𝑒_, 𝐼_𝑐𝑜𝑚𝑚𝑜𝑛_(𝑡) and 𝐼^𝑖^ (𝑡) represent the time-varying input currents either shared across the population of motor neurons or specific to the motor neuron (see *Inputs to the motor neurons*). The offset 𝐼^𝑖^ represents the minimum input required for the depolarizing current to become non-zero.
- Each presynaptic Renshaw spike induced a fixed hyperpolarizing (-5 µA) jump Δ𝐼_𝑠𝑦𝑛𝑎𝑝𝑡𝑖𝑐_ in 𝐼^𝑖^_𝑠𝑦𝑛𝑎𝑝𝑡𝑖𝑐_, which was further scaled by the Renshaw cell-to-motor neuron synaptic weight. The decay rate of this inhibitory current was governed by 𝜏^𝑖^_𝑠𝑦𝑛𝑎𝑝𝑡𝑖𝑐_, which was either manually specified or set equal to the motor neuron’s membrane time constant (𝑅^𝑖^ × 𝐶^𝑖^). Thus, 𝐼^𝑖^_𝑠𝑦𝑛𝑎𝑝𝑡𝑖𝑐_ evolved according to 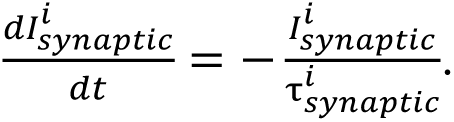

Finally, all motor neuron-specific parameters (𝐶^𝑖^, 𝑅^𝑖^, 𝜏^𝑖^𝐴𝐻𝑃, 𝐼^𝑖^𝑟ℎ𝑒𝑜𝑏𝑎𝑠𝑒) were computed from soma diameter 𝐷^𝑖^ using the power-law relations proposed by Caillet *et al.* (*62*), except in simulation scenarios where parameters were explicitly assigned. The input scaling 𝑆^𝑖^ corresponds to the normalized membrane resistance and was set to 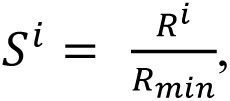 where 𝑅_𝑚𝑖𝑛_ is the membrane resistance for a soma diameter of 50 µm. Therefore, 𝑆^𝑖^ ≅ 1 for the smallest simulated motor neuron and decreased with increasing soma size. We simulated groups of 30 motor neurons with normally-distributed soma diameters (mean ± SD: 60±5 µm), consistent with smaller motor neurons (∼50–70 µm soma diameter (*63*)) typically recruited during submaximal contractions.

#### Simulated recurrent connectivity via Renshaw cells

Simulations included either a single group of motor neurons to model homonymous recurrent inhibition or two groups of motor neurons to model heteronymous recurrent inhibition between synergist muscles.

For a single simulated group of motor neurons, we instantiated 𝑛_𝑅𝑒𝑛𝑠ℎ𝑎𝑤_ = 10 Renshaw cells. A single *recurrent inhibition strength* parameter (𝜇_𝑑𝑖𝑠𝑦𝑛𝑎𝑝𝑡𝑖𝑐_) controlled the target mean disynaptic connectivity of the motor neuron-to-motor neuron weight matrix (WMN→MN). To achieve this, the mean monosynaptic connectivity of both the motor neuron-to-Renshaw cell (WMN→Renshaw) and Renshaw cell-to-motor neuron (WRenshaw→MN) weight matrices was set to the same value:

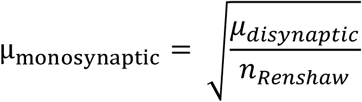

Entries of W_MN→Renshaw_ and W_Renshaw→MN_ were sampled from a normal distribution with a mean of 𝜇_𝑚𝑜𝑛𝑜𝑠𝑦𝑛𝑎𝑝𝑡𝑖𝑐_ and standard deviation of 𝜇_𝑚𝑜𝑛𝑜𝑠𝑦𝑛𝑎𝑝𝑡𝑖𝑐_ × 0.4. The effective disynaptic weights were obtained as the matrix product of the two monosynaptic weight matrices:

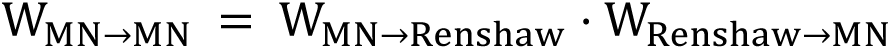

By construction the mean disynaptic connectivity between motor neurons was equal to the specified 𝜇_𝑑𝑖𝑠𝑦𝑛𝑎𝑝𝑡𝑖𝑐_ parameter, with a coefficient of variation of roughly ∼20%.

We applied the same logic when two groups of motor neurons were simulated, by defining one 𝜇_𝑑𝑖𝑠𝑦𝑛𝑎𝑝𝑡𝑖𝑐_ parameter for each ordered pair of groups (A*→*A, B→B, A→B, B→A) and instantiating 10 Renshaw cells per pair. Each set of 10 Renshaw cells was exclusively associated with a specific pair, so that each pair had its own two monosynaptic weight matrices. For example, for the A*→*B pair, the first matrix encoded the connectivity from motor neuron group A to the set of Renshaw cells A*→*B, and the second matrix encoded the connectivity from the set of Renshaw cells A*→*B to the motor neuron group B. Although such a strictly modular organization is biologically implausible, it enabled complete control of between-and within-group inhibition and thus approximated the experimentally-observed patterns of homonymous and heteronymous inhibition. Renshaw cells were modeled as leaky integrate-and-fire neurons 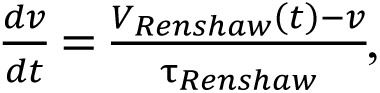 where 𝑉_𝑅𝑒𝑛𝑠ℎ𝑎𝑤_(𝑡) is the time-varying independent input to each Renshaw cell (specified as a voltage variation directly) and 𝜏_𝑅𝑒𝑛𝑠ℎ𝑎𝑤_ is the time constant, set to 8 ms (*7*). Each Renshaw cell had a resting membrane potential and reset voltage of 0 mV, a firing threshold of 10 mV, and a refractory period of 5 ms. Each Renshaw cell’s independent input 𝐼_𝑅𝑒𝑛𝑠ℎ𝑎𝑤_ (𝑡) was modelled as a zero-mean, 50 Hz low-pass-filtered Gaussian noise with a standard deviation of 3.33 mV. In addition, each spike from a pre-synaptic motor neuron produced an instantaneous depolarization Δ𝑉_𝐸𝑃𝑆𝑃_ of 6.7mV scaled by the motor neuron-to-Renshaw cell synaptic weight. Each spike from a Renshaw cell injected a decaying hyperpolarizing inhibitory current Δ𝐼_𝑠𝑦𝑛𝑎𝑝𝑡𝑖𝑐_ of-5 µA into its postsynaptic target motor neurons, scaled by the Renshaw cell-to-motor neuron synaptic weight. Unless otherwise specified, the synaptic delay was set to 5 ms in both directions (motor neurons to Renshaw cells and Renshaw cells to motor neurons).

#### Inputs to motor neurons

The simulated motor neurons within each group received a combination of external time-varying input currents. For each motor neuron *i*, the net external input 𝐼^𝑖^_𝑒𝑥𝑡𝑒𝑟𝑛𝑎𝑙_ (𝑡) consisted of three components: (i) a constant baseline excitatory current (𝐼_𝑏𝑎𝑠𝑒𝑙𝑖𝑛𝑒_), (ii) a common time-varying input (𝐼_𝑐𝑜𝑚𝑚𝑜𝑛_(𝑡)) shared among motor neurons within the same group, and (iii) an independent input (𝐼^𝑖^ (𝑡)) specific to each motor neuron. The common input was constructed as the sum of two zero-mean, band-limited Gaussian-noise processes: a low-frequency component (0–5 Hz) with a fixed amplitude (in-band standard deviation: 3.5 µA), and a higher-frequency component whose amplitude, center frequency and half-width varied across simulations. The spectral properties of the common input were chosen to reflect the well-established separation between a low-frequency component driving force modulations (*64*) and an additional higher-frequency component (*65*), consistent with observations from experimental data (*22*). The independent input to each motor neuron was a zero-mean Gaussian-noise process, low-pass filtered at 80 Hz and scaled such that its total power was three times the 0–5 Hz in-band power of the common input (*66*).

In simulations involving two groups of motor neurons, the common input for each group was first generated independently. A Cholesky-based whitening–recoloring transform was then applied to impose a specified between-groups correlation 𝜌(𝐼^𝐴^𝑐𝑜𝑚𝑚𝑜𝑛 ↔ 𝐼^𝐵^𝑐𝑜𝑚𝑚𝑜𝑛), thereby controlling the proportion of shared input while preserving each group’s common input amplitude. This step was important to simulate groups of motor neurons from anatomically defined synergist muscles, which are expected to share a certain level of common drive.

### Synchronization cross-histograms and their features

#### Constructing and interpreting the synchronization cross-histograms

We computed count-normalized synchronization cross-histograms for each motor neuron (experimental or simulated) from the *inhibition-receiver* perspective. For each neuron of interest (the *comparison* neuron), all other neurons from the homonymous pool or from a heteronymous pool served as *reference* neurons. For every spike of a *reference* neuron, we identified the closest preceding and closest following spike of the *comparison* neuron within a ±200 ms window. The corresponding delays were binned at the recording resolution (1/2048 s) to generate a pairwise cross-histogram for each *comparison-reference* pair. Each histogram was normalized to form a probability distribution (summing to 1), and these were then averaged across all *reference* neurons to yield one synchronization cross-histogram per *comparison* neuron. To suppress high-frequency noise arising from finite spike counts, each averaged distribution was convolved (zero-phase) with a normalized 16.7 ms Hanning window. The resulting smoothed synchronization cross-histogram represents the conditional probability of the neuron firing as a function of time relative to spikes from any *reference* neuron.

We pooled data across *reference* neurons to increase sample size and thereby obtain more accurate estimates. We also adopted the *inhibition-receiver* perspective, in which the motor neuron of interest is the *comparison* neuron, receiving inhibition from all *reference* neurons, rather than the *inhibition-sender* perspective in which the motor neuron of interest is the *reference* neuron, delivering inhibition to all *comparison* neurons. Although both perspectives are valid, we preferred the *inhibition-receiver* perspective because it reduced firing-rate–dependent bias at the pool level. With the *inhibition-receiver* perspective, cross-histogram width is influenced primarily by the *comparison* neuron’s own mean discharge rate, creating neuron-to-neuron variability within a pool. By contrast, the *inhibition-sender* perspective ties cross-histogram width to the pool’s average firing rate, so histograms from a given pool/simulation tend to share similar widths. Because our simulation-based inferences relied on pool-level summary statistics, the *inhibition-receiver* perspective yielded per-simulation summaries that more faithfully captured the diversity observed in the experimental data. Importantly, we verified that both perspectives resulted in qualitatively similar values for the *trough area*, *trough timing* and *peak height* features.

When constructing heteronymous synchronization cross-histograms, i.e., when the comparison motor neuron belonged to a different pool than the reference motor neurons, we selected the direction that maximized spike count per histogram. For example, because more motor units were identified in VL than in VM, using VM as the *comparison* pool and VL as the *reference* pool yielded higher per-histogram counts, and thus more reliable estimates than the converse. For the triceps surae, SOL served as the *comparison* pool and GM as the *reference* pool.

#### Curve-fitting

Each synchronization cross-histogram was fitted using a two-components model: (1) a *baseline* curve 𝐵𝑎𝑠𝑒𝑙𝑖𝑛𝑒(𝑡) representing the expected firing probability of a *comparison* neuron relative to the *reference* neurons’ spikes in the absence of recurrent inhibition and common input, and (2) a *peak-and-troughs* curve 𝑃𝑒𝑎𝑘&𝑇𝑟𝑜𝑢𝑔ℎ𝑠(𝑡) capturing the central peak and surrounding troughs. The baseline was modeled as the sum of two mirrored generalized logistic functions, 𝐺_𝐿1_(𝑡) and 𝐺_𝐿2_(𝑡), allowing flexible fitting of symmetric probability distributions with rising, plateau, and falling phases:

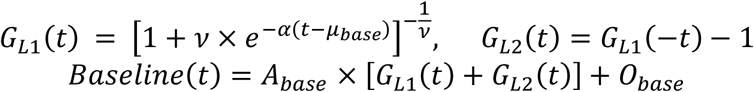

The *baseline* component included five free parameters: α, controlling the steepness of the transitions to and from the plateau; 𝜇_𝑏𝑎𝑠𝑒_, determining the horizontal centers of the logistic curves; *ν*, shaping the curve’s skew; 𝐴_𝑏𝑎𝑠𝑒_, scaling the amplitude; and 𝑂_𝑏𝑎𝑠𝑒_, providing a vertical offset.

The residual *peak-and-troughs* component was modeled by a Mexican-hat wavelet 𝑀_𝐻_(t) multiplied by a unit-height Gaussian 𝒩, as follows:

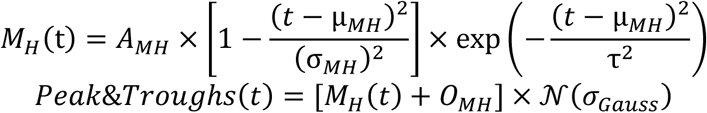

This component introduced six additional free parameters: A_𝑀𝐻_, σ_𝑀𝐻_, 𝜏, which respectively defined the amplitude, the zero-crossing of the wavelet side lobes, and the decay rate of the Mexican-hat curve; μ_𝑀𝐻_, controlling the horizontal shift of both the Mexican-hat and Gaussian; 𝜎_𝐺𝑎𝑢𝑠𝑠_, the width of the Gaussian (Gaussian center fixed at 0); and 𝑂_𝑀𝐻_, a vertical offset.

To let the *baseline* component capture most of the variance and to improve fitting convergence, we sequentially and iteratively fitted 𝐵𝑎𝑠𝑒𝑙𝑖𝑛𝑒(𝑡) and 𝑃𝑒𝑎𝑘&𝑇𝑟𝑜𝑢𝑔ℎ𝑠(𝑡) using alternating least-squares optimization, holding one component constant while fitting the other.

To ensure robust estimates, we discarded synchronization cross-histograms from analysis if any of the following rejection criteria were met: (1) fewer than 5,000 total spikes (experimental data) or 10,000 spikes (simulated data), (2) R²<0.1 for the *baseline* component fit, or (3) R²<0.75 for the composite fit. In practice, the composite fit was excellent for the vast majority of synchronization cross-histograms after exclusion (simulated data: R² = 0.97±0.04; experimental data: R² = 0.99±0.02).

#### Feature extraction

From each fitted synchronization cross-histogram, we extracted three observable features:

- *Trough area*, defined as the integral of the negative residuals (histogram minus *baseline* curve) within a predefined analysis window (see below), expressed as a percentage of total probability mass. This feature quantifies the reduction in firing probability of the *comparison* neuron relative to the baseline expectation.
- *Trough timing*, defined as the time lag with minimum autocorrelation of the residuals computed within the same analysis window as for the trough area. This feature summarizes the temporal aspect of the histogram’s troughs.
- *Peak height*, defined as the maximum value of the fitted *peak-and-troughs* curve, expressed as a probability percentage. This feature serves as a proxy for short-timescale synchrony, reflecting the higher-frequency component of the common input.

We defined the window of analysis used for the *trough area* and *trough timing* features by fitting a trapezoid to the *baseline* curve. The trapezoidal function was a piece-wise linear function, with flat outer segments at 0, a linear rise, a plateau, and a linear fall. The analysis window was defined as the post-*reference*-spike time corresponding to the plateau of the trapezoid (i.e., 0 < *t* < start of the falling edge). When computing the *trough area* backwards in time, the analysis window was shifted to the plateau’s pre-*reference*-spike interval (i.e. end of the rising edge < *t* < 0).

### Other data analyses

#### Intramuscular coherence

To assess the influence of recurrent inhibition and common input shape on the correlated activity of motor neurons in the frequency domain, we computed magnitude-squared coherence between cumulative spike trains built from simulated motor neuron activity. For each simulation, two non-overlapping random subsets of 𝑁 motor neurons were repeatedly drawn (without replacement), and two cumulative spike trains were constructed by summing the binary spike trains within each subset. The subset size 𝑁 varied from 1 up to half of the available motor neurons (here, 𝑁 ≤ 15). To keep computation time manageable, we used more iterations for smaller 𝑁 (100 at 𝑁 = 1) and progressively fewer as 𝑁 increased. Coherence between two mean-centered cumulative spike trains 𝑥 and 𝑦 at each frequency *f* was estimated with Welch’s method using 1-s Hann windows and 50% segment overlap:

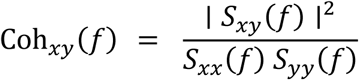

where 𝑆_𝑥𝑦_ is the cross power-spectrum and 𝑆_𝑥𝑥_, 𝑆_𝑦𝑦_ are auto power-spectra (all averaged over segments). For each 𝑁, coherence curves were averaged over random draws.

#### Recruitment thresholds

We estimated the recruitment threshold of each motor unit during the ramp-up phases of the contractions. Only valid ramps were included, defined as those in which at least 50% of samples in a 7s pre-plateau window had a z-scored derivative of the 1-Hz low-pass filtered torque exceeding 0.05. For each valid ramp, the recruitment threshold was defined as the relative torque at the time of the first spike in the earliest sequence of five consecutive spikes occurring within 1 s. For each motor unit, values were averaged across all valid ramps to obtain a single estimate.

### Experimental study

#### Participants and ethical approval

Six physically active males (age: 29.7±8.0 years, height: 177.7±3.4 cm, body mass: 73.7±5.4 kg) participated in four distinct experiments involving isometric tasks: *knee extension*, *plantarflexion*, *dorsiflexion* or *finger pinch*. The study procedures were approved by the ethics committee (CPP Ouest III; 23.00453.000166) and conducted in accordance with the Declaration of Helsinki, except for registration in a database. All participants were fully informed of the potential risks and discomforts associated before providing written informed consent.

#### Procedures

Each participant performed four distinct experiments on separate days. For the *knee extension* experiment, participants were seated on a dynamometer (Biodex System 3 Pro, Biodex Medical, USA) with the hips and right leg flexed at 80° (0° being the neutral position). For the *plantarflexion* experiment, participants sat on the same dynamometer with their hips flexed at 80° and their right leg fully extended. For the *dorsiflexion* experiment, participants were seated in a custom-built chair connected to an ankle ergometer (DinamometroGC, OT Bioelettronica, Torino, Italia). The right foot was securely attached to the footplate with the leg slightly flexed and the hips flexed at 80°. For both the plantarflexion and dorsiflexion tasks, the foot was placed perpendicular to the shank. For the *pinch* experiment, participants held a dynamometer (Cor2, OT Bioelettronica, Torino, Italia) between the radial side of the index finger and the thumb (Fig. 5).

In each experiment, the session began with a standardized warm up, followed by three maximal voluntary contractions (MVC) for 3 s, with 60 s of rest in between. Peak MVC force was considered as the maximal value obtained from a moving average window of 250 ms. Participants then performed 10 trapezoidal isometric contractions, each consisting of a 5-s ramp followed by a 20-s plateau at 10% MVC, with 30-s of rest between contractions. This was followed by 20 trapezoidal isometric contractions, each with a 5-s ramp and 10-s plateau at 40% MVC, with at least 30-s of rest in between. We instructed the participants to follow the ramp on the first five contractions only. This large number of contractions ensured the acquisition of sufficient motor unit firings to allow for a robust assessment of recurrent inhibition.

#### Surface electromyography recordings

Electromyographic (EMG) signals were recorded using adhesive grids of 64 electrodes, with an inter-electrode distance (IED) of 8 mm or 4 mm (GR08MM1305 or GR04MM1305, OT Bioelettronica, Italy) (Fig. 5A). In the *knee extension* experiment, four grids (8-mm IED) were placed over the vastus lateralis (VL), and two grids (4-mm IED) were placed over the vastus medialis (VM). In the *plantarflexion* experiment, four grids (4-mm IED) were placed over the Gastrocnemius medialis (GM), and two grids (4-mm IED) were placed over Soleus (SOL), targeting its medial and lateral portion. In the *dorsiflexion* experiment, four grids (4-mm IED) were placed over the Tibialis anterior (TA). Finally, in the *pinch* experiment, one grid (4-mm IED) was placed over the first dorsal interosseous (FDI) muscle. When possible, multiple grids were placed over the same muscle to maximize the number of decomposed motor units.

Prior to electrode placement, the skin was shaved and cleaned with an abrasive paste (Nuprep, Weaver and Company, USA). The adhesive grids were secured with double-sided adhesive foam layers (SpesMedica, Battipaglia, Italy). The skin-electrode contact was made by filling the cavities of the adhesive layers with conductive paste (SpesMedica, Battipaglia, Italy). A reference electrode (5×5 cm; Kendall Medi-Trace; Covidien, Ireland) was placed over a bony prominence (tibia, patella, or wrist), and a strap electrode dampened with water (ground electrode) was placed around the left ankle or the wrist. The EMG signals were recorded in a monopolar montage, bandpass filtered (10–500 Hz), and digitized at a sampling rate of 2048 Hz using a multichannel acquisition system (EMG-Quattrocento; 400-channel EMG amplifier; OT Biolelettronica, Italy).

#### Decoding of motor unit firing activity

The EMG signals were decomposed using convolutive blind source separation (*68*), as implemented in the MUedit software (*69*). Prior to automatic decomposition, all channels were visually inspected, and those with low signal-to-noise ratio or artifacts were discarded. The remaining EMG signals were extended and whitened, after which fast independent component analysis was performed to retrieve the motor unit spike trains. The resulting spike trains were manually edited according to established procedures (*70*). Manual editing involved removing detected peaks that result in erroneous firing rates (outliers) and adding missed firing times that are clearly distinguishable from the noise. The motor unit pulse trains were ultimately recalculated with updated separation vectors and accepted by the operator once all the putative firing times were selected. This procedure was demonstrated to be highly reliable across operators (*71*).

Decomposition was performed independently for each electrode grid. Because the same motor units could be identified across multiple grids covering the same muscle, duplicate units were detected by comparing their spike trains. Firing times that occurred within 0.5 ms interval were considered as common between motor units; and motor units that shared more than 30% of their firing times were considered as duplicates (*69*). When duplicated motor units were identified, only the motor unit with the lowest coefficient of variation of its inter-spike intervals was retained.

Prior to analyses, the spike trains were realigned based on the onset of their average motor unit action potential (MUAP). The average MUAP shape was calculated using spike-trigger averaging, and its onset was defined as the time at which it first crossed a threshold set at six standard deviations from the baseline (*72*).

### Simulation-based inference

We performed simulation-based inference using the sbi Python toolbox (*73*) with sequential neural *posterior* estimation (SNPE (*21*)) to infer physiological parameters from observed features derived from motor neuron spiking activity. This approach involved the following steps: (1) specifying *priors* over a subset of parameters relevant to our research question, (2) simulating spike trains with our simulation model, (3) reducing the output of each simulation to a set of summary statistics derived from observable features, and (4) training a neural density estimator to approximate the *posterior* distribution (i.e., the probability of parameter values given the observed features). This framework was validated from both experimental and simulated data.

### Observable features and inferred parameters

Each simulated and experimentally recorded motor unit spike train was reduced to four observable features: *trough area*, *trough timing*, *peak height*, and mean firing rate. Note that the inference procedure was applied independently for each muscle and contraction intensity, at the level of the group of identified motor neurons. Specifically, motor units were pooled across participants, and each observable feature was characterized by its mean, median, standard deviation, and interquartile range. These summaries yielded a 16-elements feature-summary vector (4 observables × 4 summary statistics) per experimental condition, which the neural density estimator mapped to a *posterior* distribution over physiological parameters. In the within-muscle case, we inferred five physiological parameters: (1) baseline excitation (common input offset 𝐼_𝑏𝑎𝑠𝑒𝑙𝑖𝑛𝑒_), (2) recurrent inhibition strength (mean disynaptic connectivity 𝜇_𝑑𝑖𝑠𝑦𝑛𝑎𝑝𝑡𝑖𝑐_), (3) amplitude of the higher-frequency component of the common input σ(𝐼_𝑐𝑜𝑚𝑚𝑜𝑛,ℎ𝑖𝑔ℎ_), and the (4) center frequency 𝑓_𝑐𝑒𝑛𝑡𝑒𝑟_ and (5) half-width 𝑓_ℎ𝑎𝑙𝑓−𝑤𝑖𝑑𝑡ℎ_ of that higher-frequency component. In the between-muscles case, we inferred heteronymous recurrent inhibition strength in both directions, i.e., (1) from muscle A to muscle B (𝜇^𝐴→𝐵^𝑑𝑖𝑠𝑦𝑛𝑎𝑝𝑡𝑖𝑐) and (2) from muscle B to muscle A (𝜇^𝐵→𝐴^𝑑𝑖𝑠𝑦𝑛𝑎𝑝𝑡𝑖𝑐 ), and (3) the proportion of shared common input between muscles 𝜌(𝐼^𝐴^𝑐𝑜𝑚𝑚𝑜𝑛 ↔ 𝐼^𝐵^𝑐𝑜𝑚𝑚𝑜𝑛). Of note, *posterior* estimates of heteronymous recurrent inhibition strength were highly similar in both directions (e.g., A→B inhibition ≅ B→A inhibition). For clarity, we therefore report a single bidirectional value per synergist muscles pair in the Results section. Because the proportion of common input was weakly identifiable (uncertain marginal *posterior*), it is not reported.

#### *Priors* and training simulations

The physiological parameters to infer were chosen to estimate recurrent inhibition and higher-frequency common input while keeping the dimensionality of the parameter space tractable. We selected *priors* ranges broad enough for the simulated feature summaries to span the entire range of feature summaries derived from experimental data (Fig S1), yet constrained enough to avoid wasting simulation resources on unobserved regions of the parameter space. In the within-muscle case, independent uniform *priors* were used for all five parameters (𝐼_𝑏𝑎𝑠𝑒𝑙𝑖𝑛𝑒_, 𝜇_𝑑𝑖𝑠𝑦𝑛𝑎𝑝𝑡𝑖𝑐_, σ(𝐼_𝑐𝑜𝑚𝑚𝑜𝑛,ℎ𝑖𝑔ℎ_), 𝑓_𝑐𝑒𝑛𝑡𝑒𝑟_, 𝑓_ℎ𝑎𝑙𝑓−𝑤𝑖𝑑𝑡ℎ_; bounds in Table 1). We generated 12,000 simulations as training data, yielding one feature-summary vector per simulation. For the between-muscles case, parameters used the inferred within-muscle *posteriors* as informed *priors*, while between-muscles parameters (𝜇^𝐴→𝐵^𝑑𝑖𝑠𝑦𝑛𝑎𝑝𝑡𝑖𝑐, 𝜇^𝐵→𝐴^𝑑𝑖𝑠𝑦𝑛𝑎𝑝𝑡𝑖𝑐, 𝜌(𝐼^𝐴^𝑐𝑜𝑚𝑚𝑜𝑛 ↔ 𝐼^𝐵^𝑐𝑜𝑚𝑚𝑜𝑛)) used independent uniform *priors* (bounds in Table 1). This approach concentrated simulation effort where within-muscle parameters were most probable while allowing between-muscle parameters to vary freely. We generated 20,000 simulations to train the neural density estimator in the between-muscles case: for each synergist pair (VL↔VM, GM↔SOL) and intensity (10% and 40% MVC), 5,000 parameter sets were drawn, with within-muscle values sampled from the corresponding *posterior* and between-muscles values from their uniform *prior*.

**Table 1.**
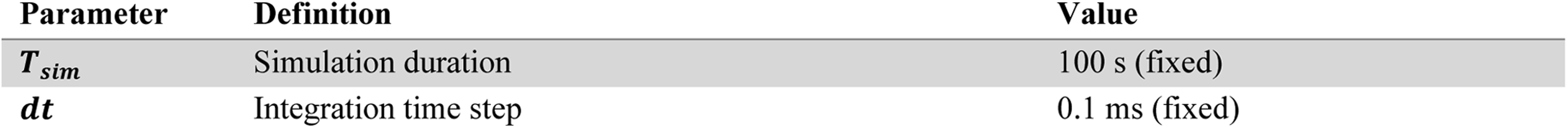
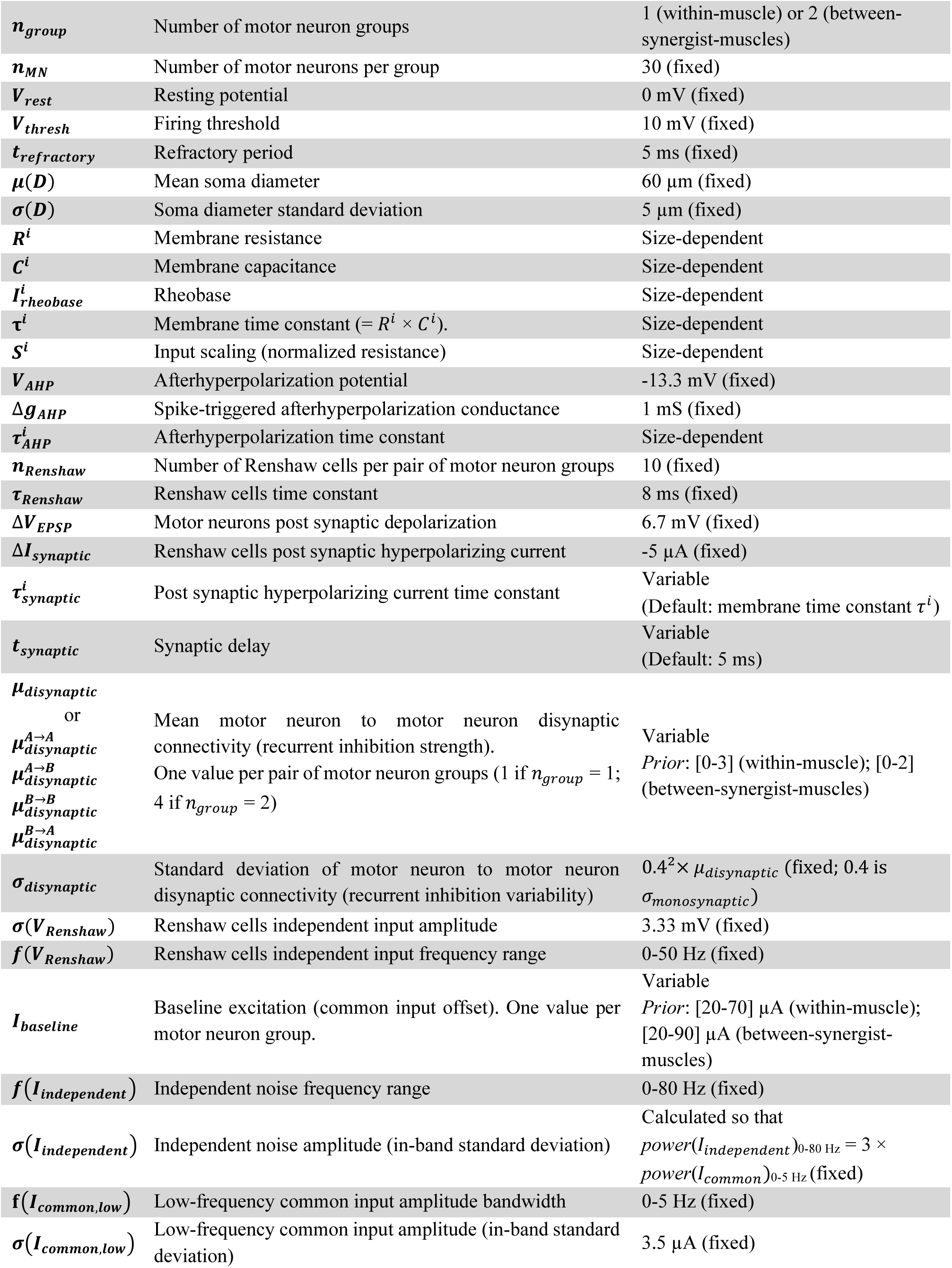
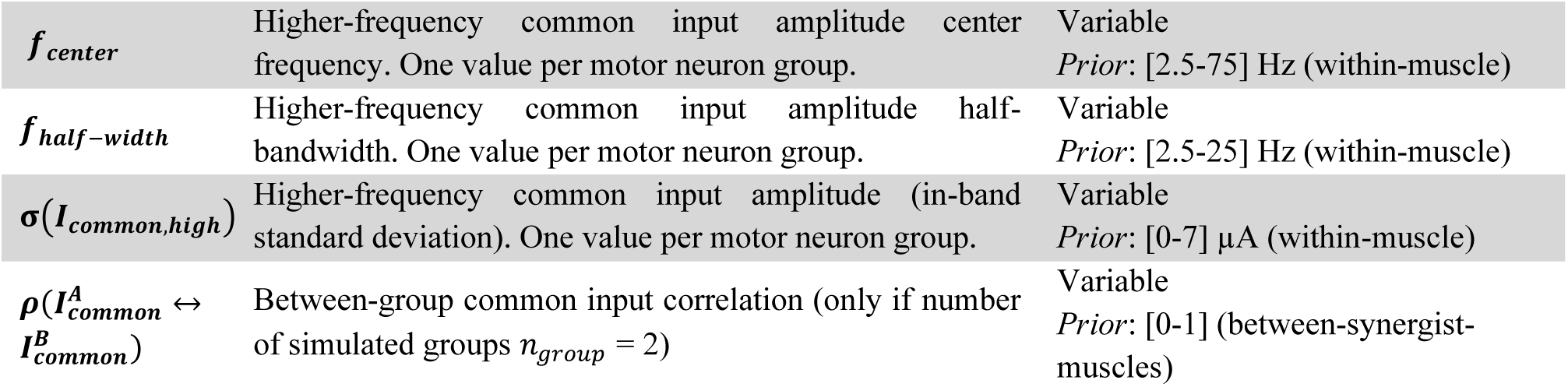
Simulation parameters.

| Parameter | Definition | Value |
| --- | --- | --- |
| $T_{sim}$ | Simulation duration | 100 s (fixed) |
| $dt$ | Integration time step | 0.1 ms (fixed) |
| $n_{group}$ | Number of motor neuron groups | 1 (within-muscle) or 2 (between-synergist-muscles) |
| $n_{MN}$ | Number of motor neurons per group | 30 (fixed) |
| $V_{rest}$ | Resting potential | 0 mV (fixed) |
| $V_{thresh}$ | Firing threshold | 10 mV (fixed) |
| $t_{refractory}$ | Refractory period | 5 ms (fixed) |
| $\mu(D)$ | Mean soma diameter | 60 $\mu\text{m}$ (fixed) |
| $\sigma(D)$ | Soma diameter standard deviation | 5 $\mu\text{m}$ (fixed) |
| $R^i$ | Membrane resistance | Size-dependent |
| $C^i$ | Membrane capacitance | Size-dependent |
| $I_{rheobase}^i$ | Rheobase | Size-dependent |
| $\tau^i$ | Membrane time constant ($= R^i \times C^i$). | Size-dependent |
| $S^i$ | Input scaling (normalized resistance) | Size-dependent |
| $V_{AHP}$ | Afterhyperpolarization potential | -13.3 mV (fixed) |
| $\Delta g_{AHP}$ | Spike-triggered afterhyperpolarization conductance | 1 mS (fixed) |
| $\tau_{AHP}^i$ | Afterhyperpolarization time constant | Size-dependent |
| $n_{Renshaw}$ | Number of Renshaw cells per pair of motor neuron groups | 10 (fixed) |
| $\tau_{Renshaw}$ | Renshaw cells time constant | 8 ms (fixed) |
| $\Delta V_{EPSP}$ | Motor neurons post synaptic depolarization | 6.7 mV (fixed) |
| $\Delta I_{synaptic}$ | Renshaw cells post synaptic hyperpolarizing current | -5 $\mu\text{A}$ (fixed) |
| $\tau_{synaptic}^i$ | Post synaptic hyperpolarizing current time constant | Variable<br>(Default: membrane time constant $\tau^i$) |
| $t_{synaptic}$ | Synaptic delay | Variable<br>(Default: 5 ms) |
| $\mu_{disynaptic}$<br>or<br>$\mu_{disynaptic}^{A \rightarrow A}$<br>$\mu_{disynaptic}^{A \rightarrow B}$<br>$\mu_{disynaptic}^{B \rightarrow B}$<br>$\mu_{disynaptic}^{B \rightarrow A}$ | Mean motor neuron to motor neuron disynaptic connectivity (recurrent inhibition strength).<br>One value per pair of motor neuron groups (1 if $n_{group} = 1$; 4 if $n_{group} = 2$) | Variable<br>*Prior*: [0-3] (within-muscle); [0-2] (between-synergist-muscles) |
| $\sigma_{disynaptic}$ | Standard deviation of motor neuron to motor neuron disynaptic connectivity (recurrent inhibition variability) | $0.4^2 \times \mu_{disynaptic}$ (fixed; 0.4 is $\sigma_{monosynaptic}$) |
| $\sigma(V_{Renshaw})$ | Renshaw cells independent input amplitude | 3.33 mV (fixed) |
| $f(V_{Renshaw})$ | Renshaw cells independent input frequency range | 0-50 Hz (fixed) |
| $I_{baseline}$ | Baseline excitation (common input offset). One value per motor neuron group. | Variable<br>*Prior*: [20-70] $\mu\text{A}$ (within-muscle); [20-90] $\mu\text{A}$ (between-synergist-muscles) |
| $f(I_{independent})$ | Independent noise frequency range | 0-80 Hz (fixed) |
| $\sigma(I_{independent})$ | Independent noise amplitude (in-band standard deviation) | Calculated so that<br>$power(I_{independent})_{0-80 \text{ Hz}} = 3 \times power(I_{common})_{0-5 \text{ Hz}}$ (fixed) |
| $f(I_{common,low})$ | Low-frequency common input amplitude bandwidth | 0-5 Hz (fixed) |
| $\sigma(I_{common,low})$ | Low-frequency common input amplitude (in-band standard deviation) | 3.5 $\mu\text{A}$ (fixed) |
| $f_{center}$ | Higher-frequency common input amplitude center frequency. One value per motor neuron group. | Variable<br>Prior: [2.5-75] Hz (within-muscle) |
| $f_{half-width}$ | Higher-frequency common input amplitude half-bandwidth. One value per motor neuron group. | Variable<br>Prior: [2.5-25] Hz (within-muscle) |
| $\sigma(I_{common,high})$ | Higher-frequency common input amplitude (in-band standard deviation). One value per motor neuron group. | Variable<br>Prior: [0-7] μA (within-muscle) |
| $\rho(I_{common}^A \leftrightarrow I_{common}^B)$ | Between-group common input correlation (only if number of simulated groups $n_{group} = 2$) | Variable<br>Prior: [0-1] (between-synergist-muscles) |

#### Neural density estimator

We used the sbi toolbox default SNPE implementation with a masked autoregressive flow (MAF (*74*)) as the neural density estimator. Briefly, SNPE trains the neural density estimator to maximize the log-probability-density of the ground-truth parameters under the learned *posterior*, and MAF is a normalizing flow network that learns to map a simple base probability distribution to a target probability distribution (the *posterior* in this case). We kept the default architecture and preprocessing (50 hidden features, 5 transforms, independent z-scoring of parameters and features, and a standard-normal base distribution). We performed stochastic gradient descent with a training batch size of 200, a learning rate of 5×10⁻⁴, a 10% validation split, and early stopping after 20 epochs without improvement (explicit cap of 2,000 epochs). We trained separate estimators for the within-muscle and between-muscle conditions. We repeated the training of each estimator with different seeds to assess dependence on random initialization: the inferred *posteriors* remained qualitatively stable under different initial conditions; variability was largest for conditions with broad *posteriors* (i.e., high uncertainty, like for SOL at 40% MVC). *Posteriors* reported in the Results section are derived from a representative training run.

#### Posterior sampling

For each observed feature-summary vector, *posterior* samples were drawn from the trained neural density estimator using the default *posterior*-sampling procedure in the *sbi* toolbox. Because the density estimator is trained under bounded *priors*, samples falling outside the *prior* support were automatically rejected. In this context, high rejection rates indicate that the simulation model and chosen *priors* provide poor support for the observed data. In our experimental data, *posterior* rejection rates remained low, consistent with the fact that the observed feature summaries were well covered by the *prior*-predictive distribution of the simulated data (Fig. S1). For the additional sensitivity analysis on simulated data examining the relationship between synchronization cross-histogram mean spike count and inference usability (Fig. S5 C-D), we considered a *posterior* to be usable when at least 5% of samples fell within the *prior* support.

#### Validation

To assess the estimator’s ability to recover parameters under correct-model assumptions, we withheld 10% of simulations from training and evaluated the *posteriors* estimated from the trained estimator on this held-out dataset. Values for each parameter were mapped to a [0,1] scale relative to *prior* bounds. As recommended by the literature (*25*, *26*), we assessed *accuracy* (how close the estimator’s marginal *posteriors* are to the held-out ground truths; Supplementary Fig. S2A) and *calibration* (how well the *posterior* uncertainty matches the frequency with which the ground truth is contained; Supplementary Fig. S2B) for each parameter. Accuracy was quantified by calculating RMSE and Pearson’s correlation coefficient *r* between the ground-truth values across held-out simulations and the corresponding *posterior* modes (the parameter values with highest *posterior* probability). RMSE captures absolute errors on the [0,1] scale, whereas *r* captures how well the estimator preserves the pattern of variation across simulations, irrespective of any bias. As a reference, we also computed the RMSE and Pearson’s *r* expected from the *prior* (independent samples from a uniform distribution over [0,1]). Calibration was assessed using coverage curves. For each parameter and each held-out simulation, we calculated the highest-*posterior*-density intervals for percentages of the *posterior* probability masses between 0% (interval of zero length) and 100% (the full [0,1] range). At each percentage, we then measured the proportion of ground-truth values that fell inside the corresponding highest-*posterior*-density interval across held-out simulations. A perfectly calibrated estimator yields a coverage curve that follows the identity line; for example, 50% highest-*posterior*-density intervals contain the ground truth in 50% of simulations. Overconfidence and underconfidence are reflected in curves lying below or above this line, respectively. We summarized miscalibration for each parameter by the mean absolute deviation between its coverage curve and the identity line.

In addition, we performed *posterior* predictive checks by testing whether simulations whose parameters came from the inferred *posteriors* accurately reproduced the experimental data. For each condition we drew 100 parameter samples from the corresponding *posterior* and used them as inputs to the mechanistic model to simulate spike trains. For each simulated spike train, we computed the four observable features (*trough area*, *peak height*, *trough timing*, and mean firing rate) and then averaged each feature across motor neurons, yielding a distribution of 100 *posterior*-predicted feature means per condition. We then standardized (mean=0, standard deviation=1) the values for each feature relative to the experimentally observed per-condition means. Finally, for each feature and across conditions, we computed the RMSE and Pearson’s *r* between the *posterior*-predicted feature means (center of mass of each condition’s *posterior*-predictive distribution) and the experimentally observed feature means.

#### Robustness to model misspecification

Our model assumes a single common input to the motor neuron pool, which is a reasonable approximation for most pools (*64*). To assess the sensitivity of the inference to the spatial structure of inputs, we generated 500 additional simulations from the same *priors* as in the main analysis, in which two common inputs projected heterogeneously to the motor neuron pool. The weight assigned to each motor neuron was drawn from a uniform distribution between 0 and 1, with the constraint that the two weights summed to 1 and had an expected mean weight of 0.5 across motor neurons for both inputs. With this design, the only difference from the original simulations was the spatial distribution and dimensionality of the inputs. We then applied the same neural density estimator, trained on the original simulations, to this new synthetic dataset. We evaluated inference accuracy by comparing *posterior* modes to ground truth. Estimates of recurrent inhibition strength were largely unaffected (Pearson’s *r* = 0.88 for two common inputs *vs* 𝑟 = 0.89 for one common input; RMSE on [0,1] normalized values: 0.16 *vs* 0.15). However, the amplitude of the higher-frequency common input tended to be underestimated in the presence of two inputs, resulting in reduced accuracy (RMSE: 0.25 *vs* 0.15). Importantly, a high correlation with ground truth was preserved (Pearson’s *r* = 0.75 *vs r* = 0.85), indicating that relative comparisons across muscles or intensities remain informative.

We further assessed the robustness to model misspecification by examining the influence of the number of simulated motor neurons. Specifically, we generated simulations with 100 motor neurons and 30 Renshaw cells. Inference was then performed using either the full set of motor neurons or random subsets of 10, 20, or 30 motor neurons, applying the neural density estimator trained on the original simulations (30 motor neurons, 10 Renshaw cells). To maintain comparable levels of inhibition, we adjusted the *prior* range of recurrent inhibition strength to [0, 0.9], instead of the [0, 3] range used in the original simulations, as the net inhibitory effect depends on the total number of simulated motor neurons. All other parameters were unchanged. Despite this deliberate misspecification, inferred recurrent inhibition strength, evaluated using *posterior* modes, remained well correlated with ground truth across simulations (mean Pearson’s *r* = 0.74).

We also tested the sensitivity of our approach to spike count, as determined by the number of identified motor neurons, recording duration, and firing rates. To this end, we simulated data with varying recording durations (20, 50, 100, and 200 s) and analyzed subsets of 10, 20, 30, or 100 motor neurons drawn from a full set of 100 simulated neurons. The accuracy of the inferred recurrent inhibition strength remained stable across combinations of motor neuron sample sizes and recording durations (Fig. S5A). However, this result should be interpreted with caution, as shorter recordings or smaller motor neuron samples substantially increased the probability that a condition was excluded due to failure to meet quality-control criteria, for example because too few reliable synchronization cross-histograms were available to compute the feature summary statistics (Fig. S5 B). Together, these results indicate that recordings with smaller spike count are less likely to satisfy the criteria for inference, but when the data pass quality control, the inferred recurrent inhibition strength is largely insensitive to spike count. An additional analysis performed on simulated data showed that a simulation run had a ≥95% probability of being accepted for inference when the mean spike count per synchronization cross-histogram (averaged across all cross-histograms in that simulation run) was approximately 15,000 (Fig. S5C).

## Author Contributions

Conceptualization: FD, FH, SA, DF Investigation: FD, FH

Formal analysis, methodology & visualization: FD Supervision: FH, SA, TC

Writing - original draft: FD, FH

Writing – review & editing: FD, FH, SA, TC, DF

## Data and materials availability

All data are available in the main text or the supplementary materials. All code used for data analysis and modeling is available at: https://github.com/FrancoisDernoncourt/Mapping_Recurrent_Inhibition

The full dataset (raw electromyography data and edited motor unit spike trains) is available at: https://figshare.com/s/c4da3eb57c88a9b07ddc

## Supplementary materials

**Figure S1.**
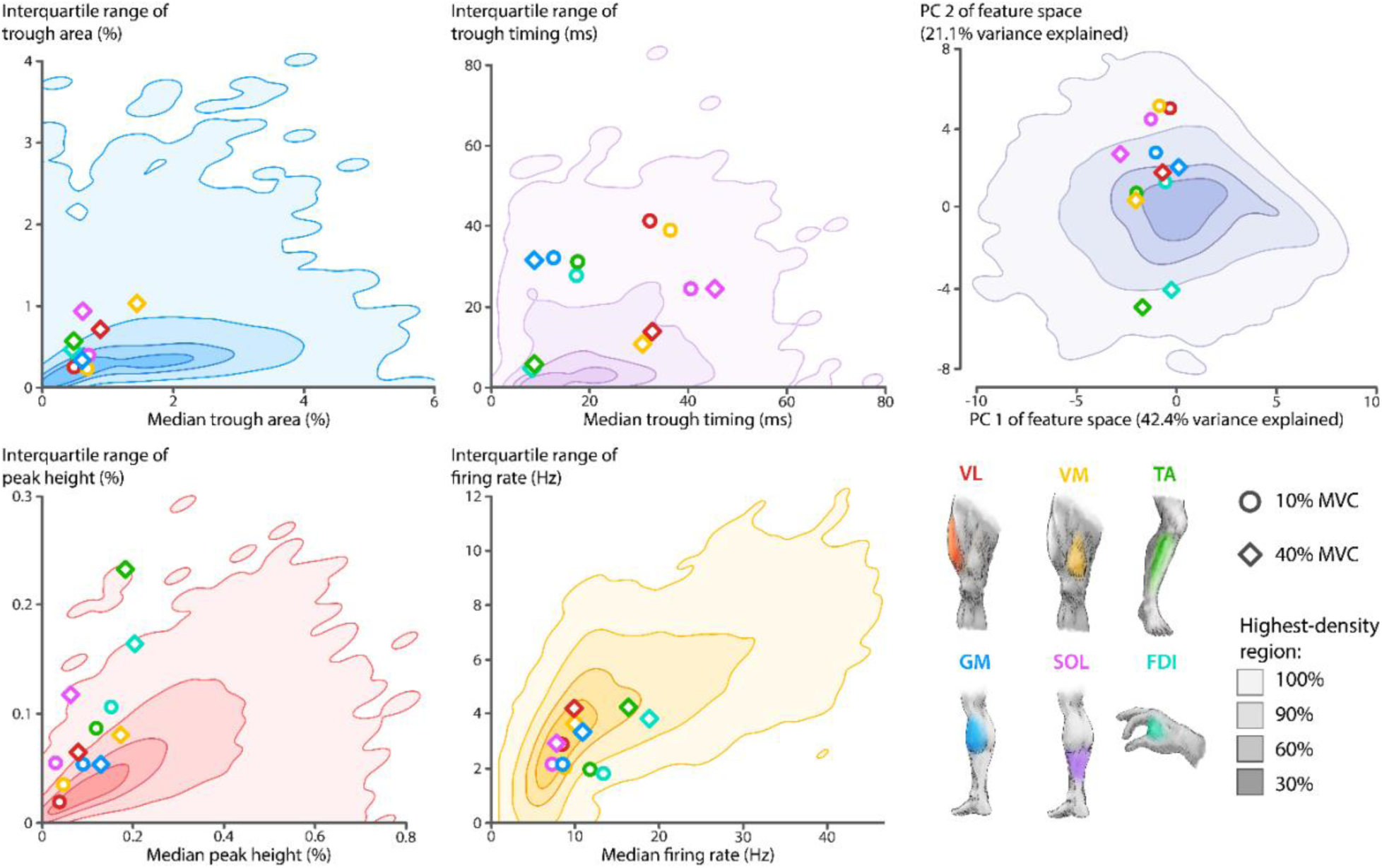
***Coverage of experimental observations by the simulated training data.*** Shaded contours indicate the highest-density regions of the training simulations used to train the neural density estimator, and markers indicate data from experimental conditions. The four left panels show the median and interquartile range of trough area, trough timing, peak height, and mean firing rate. They demonstrate that experimental observations fall within the feature distribution generated by the simulations. The top right panel shows a two-dimensional principal component representation of the full summary-statistics feature space (4 observed features × 4 summary statistics). Whereas the other panels show that the experimental observations are covered for each feature considered separately, the principal component representation shows that they are also covered when all features are considered jointly. VL: Vastus lateralis; VM: Vastus medialis; GM: Gastrocnemius medialis; SOL: Soleus; FDI: First dorsal interosseous, TA: Tibialis anterior; MVC: Maximal voluntary contraction.

**Figure S2.**
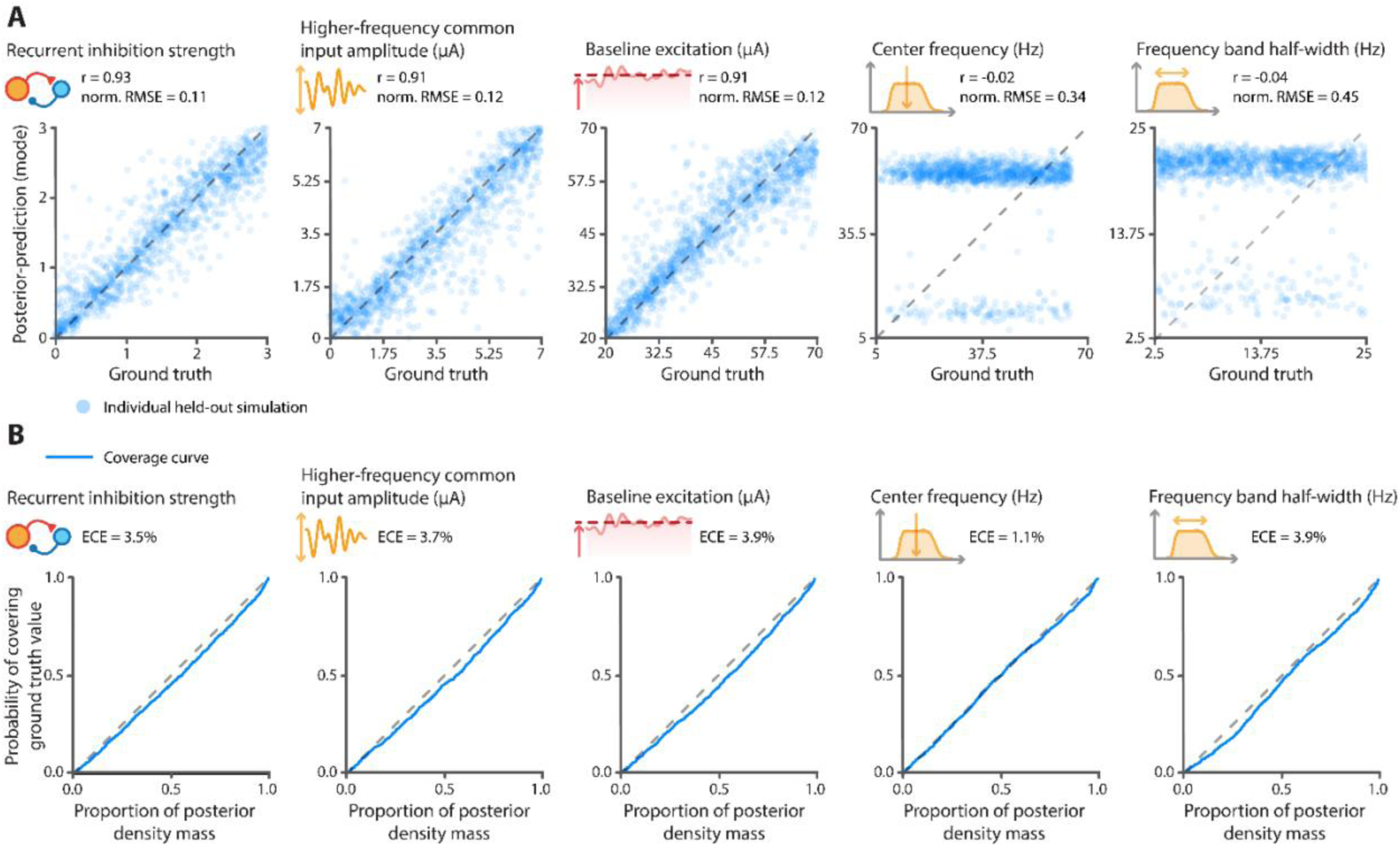
Validation of inference on held-out simulated data. **(A)** Accuracy of parameter recovery. The panels show, for each parameter, the relationship between the simulated ground-truth (x-axis) and the corresponding *posterior*-predicted mode (y-axis). Norm. RMSE: root mean square error for the [0,1]-normalized parameters. Under a uniform *prior*, expected normalized RMSE ≅ 0.41, and expected Pearson’s r = 0. **(B)** Calibration of *posterior* uncertainty. Each panel shows the calibration of the neural density estimator for one parameter. The expected calibration error (ECE) is the average deviation from the identity line, i.e., perfect calibration.

**Figure S3.**
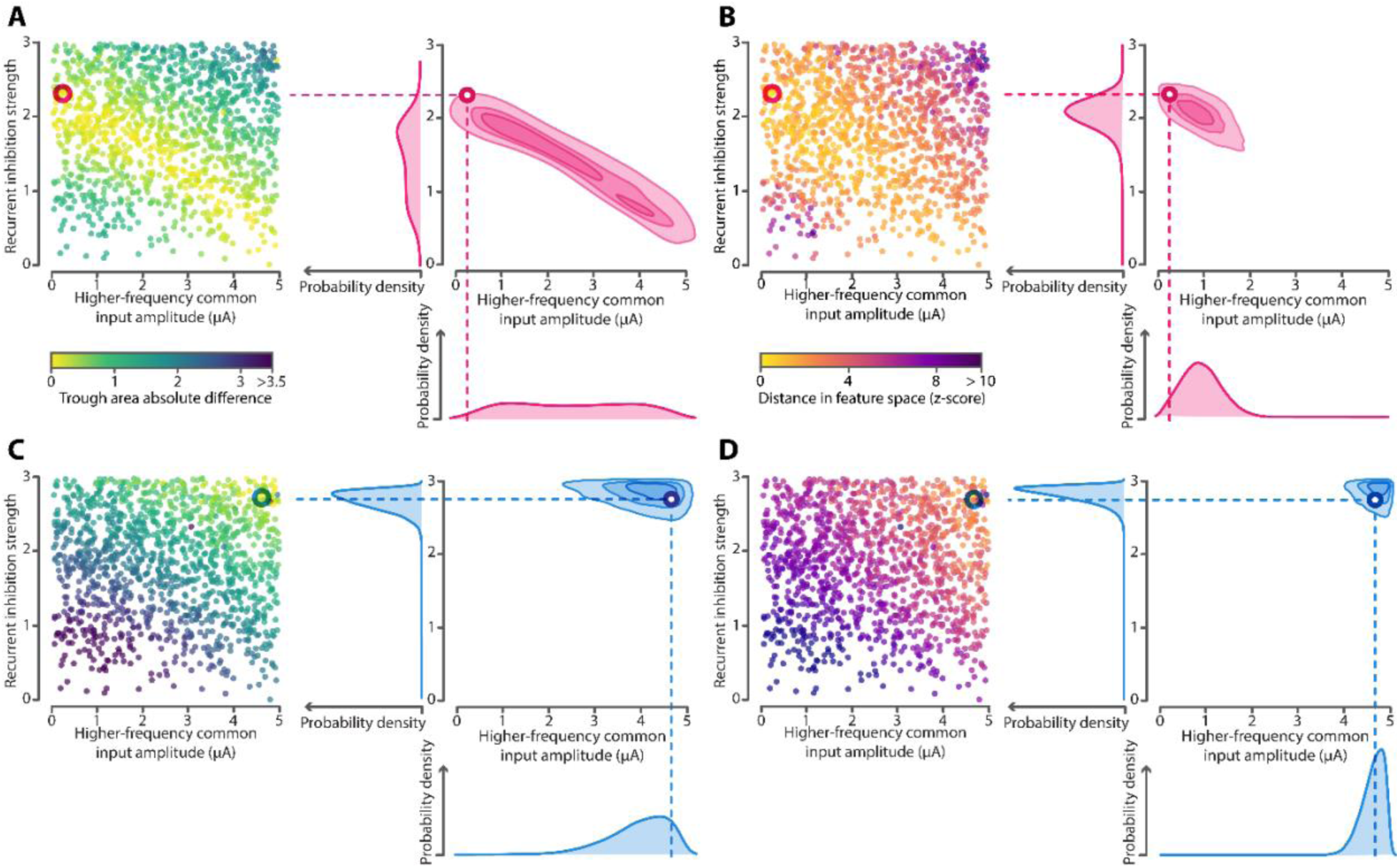
*Posteriors* in ambiguous *vs* unambiguous simulation examples. **(A)** These figures are based on the same simulated dataset as in Fig. 3A-B. Each dot represents a simulation, with color indicating the absolute difference in mean trough area relative to the selected simulation example (red circle). The right panel shows the corresponding joint posterior when inference is conditioned on the mean trough area alone. The resulting broad ridge-shaped *posterior* indicates strong indeterminacy. **(B)** The same data as in panel A, but colors now indicate the z-score–normalized Euclidean distance to the selected simulation (same as in Panel A) in the full summary-statistics feature space (4 observed features × 4 summary statistics), and inference is conditioned on this full feature set. The resulting *posterior* is more constrained than in panel A. **(C)** Same as panel A, but for a second selected simulation example located in a less ambiguous region of the parameter space. **(D)** Same as panel B, but for the simulation example shown in C. For this second example, uncertainty is already reduced in panel C and becomes very low in panel D.

**Figure S4.**
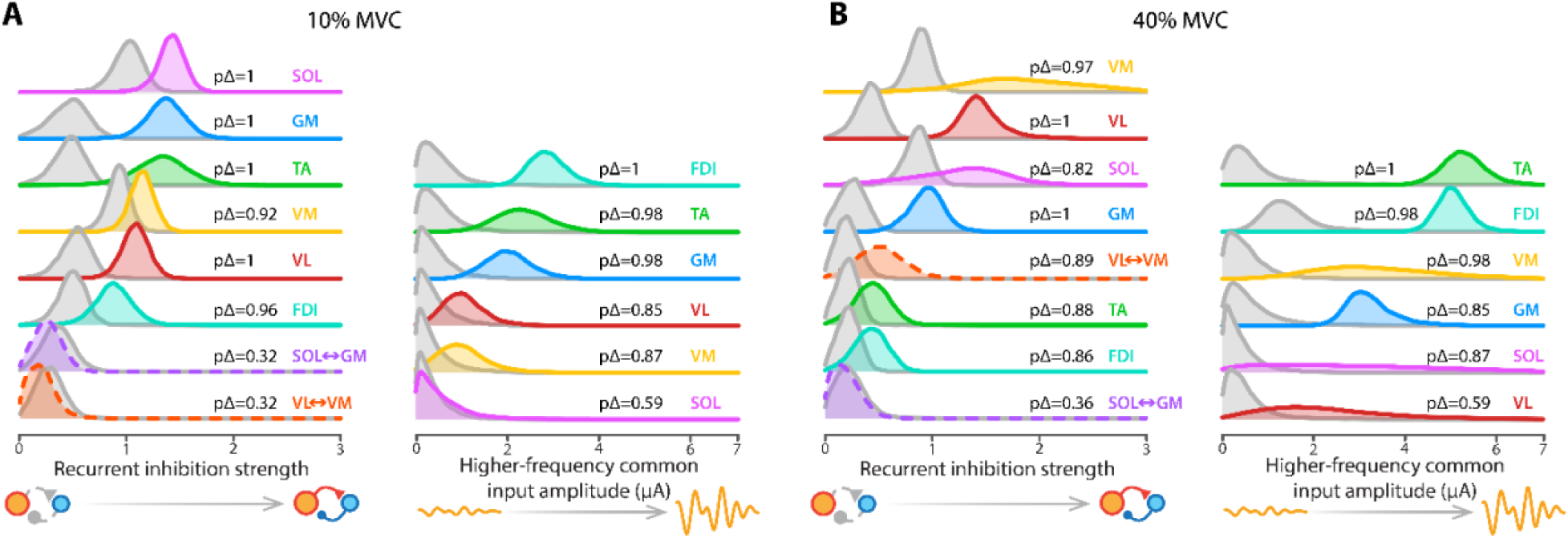
Comparison of *posteriors* inferred from original and shuffled experimental data. **(A)** Marginal *posterior* distributions of recurrent inhibition strength and higher-frequency common input amplitude are shown for each muscle and muscle pair at 10% MVC. Colored distributions correspond to inference from the original experimental data whereas gray distributions were obtained after independently circularly shifting each motor unit spike train. pΔ values indicate the probability that the parameter inferred from the original data exceeds that inferred from the shuffled data. **(B)** Same as in panel A for 40% MVC. Note that slight differences between the *posteriors* shown here and those shown in Fig. 6 are due to variability in *posterior* sampling from the same trained neural density estimator. VL: Vastus lateralis; VM: Vastus medialis; GM: Gastrocnemius medialis; SOL: Soleus; FDI: First dorsal interosseous, TA: Tibialis anterior; MVC: Maximal voluntary contraction.

**Figure S5.**
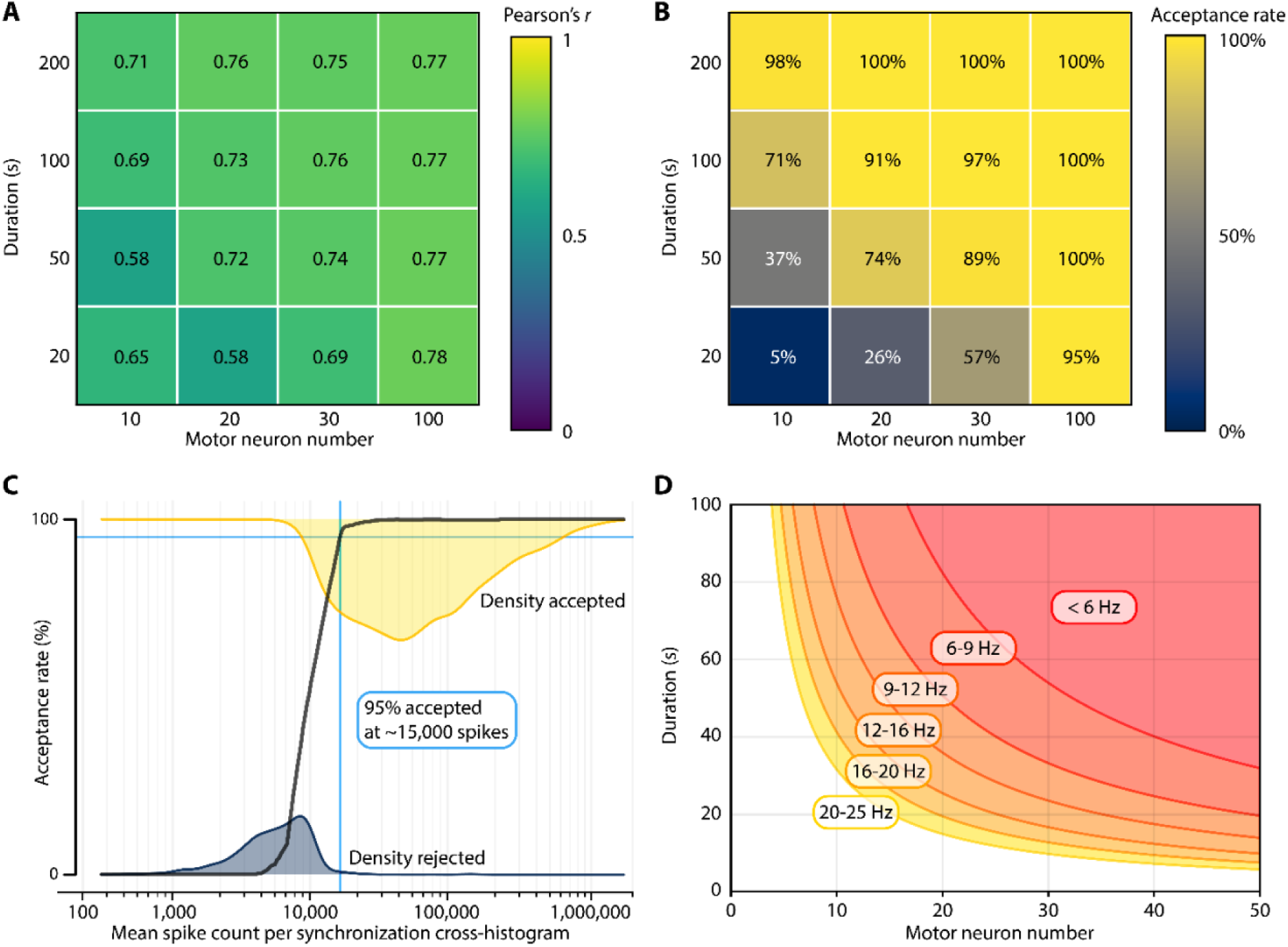
Sensitivity of the simulation-based inference pipeline to recording duration, motor neuron number, and spike count. **(A)** Mean Pearson’s *r* between ground-truth recurrent inhibition strength and its inferred value (*posterior* mode) across simulations is shown for different recording durations and numbers of sampled motor neurons. Only simulations that passed the quality-control criteria (see Methods) are included. **(B)** Percentage of simulations retained for inference across different recording durations and numbers of sampled motor neurons. **(C)** Relationship between the mean spike count per synchronization cross-histogram and the acceptance rate for inference. Yellow and dark blue shaded areas show the densities of accepted and rejected simulations, respectively, and the black curve shows the corresponding acceptance rate. In this simulation set, an average of ∼15,000 spikes per synchronization cross-histogram was associated with a 95% acceptance rate, i.e. a 95% probability that the quality-control criteria are met. **(D)** Practical guideline derived from the same simulations, relating recording duration (y-axis), motor neuron number (x-axis) and mean firing rate (color) associated with a ≥95% probability of passing the quality-control criteria.

