## Supplementary figures for "Probabilistic inference of Homonymous and Heteronymous Recurrent Inhibition in Human Muscles from Large-Scale Motor Neuron Recordings"

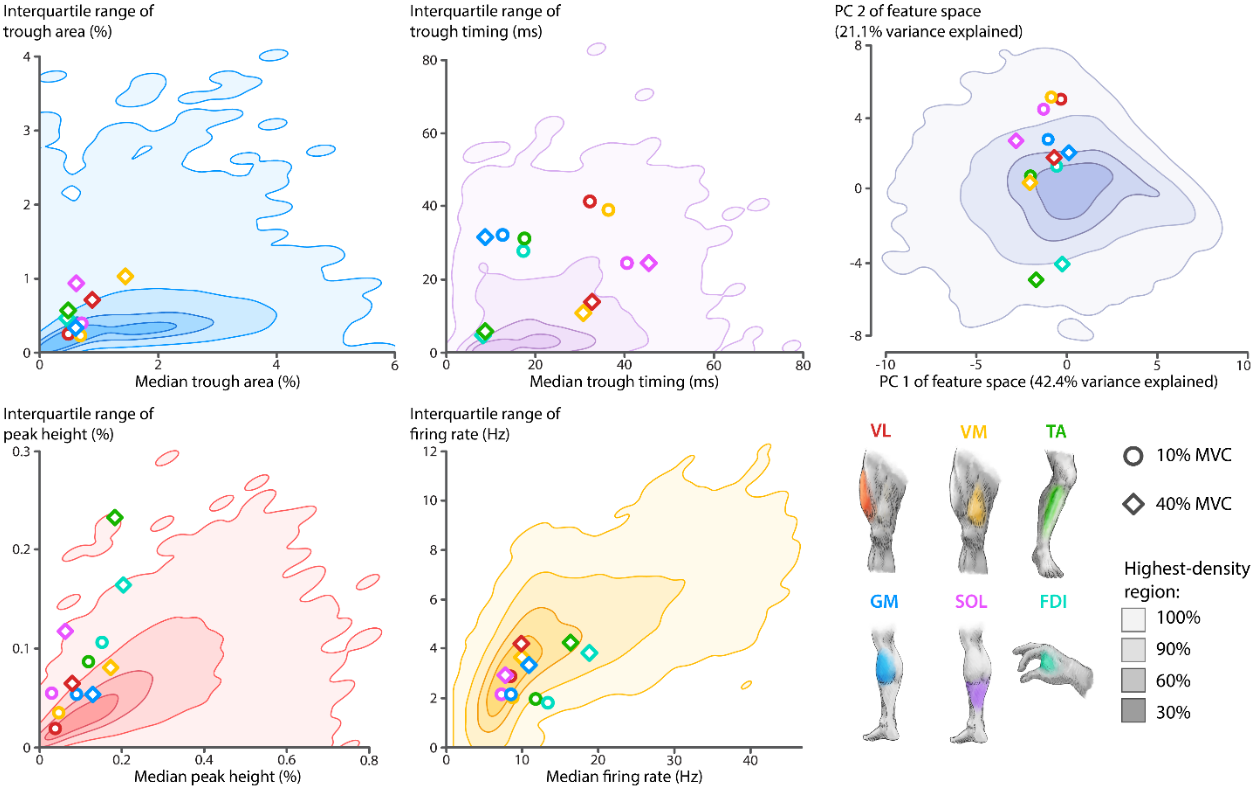


**Figure S1. Coverage of experimental observations by the simulated training data.** Shaded contours indicate the highest-density regions of the training simulations used to train the neural density estimator, and markers indicate data from experimental conditions. The four left panels show the median and interquartile range of trough area, trough timing, peak height, and mean firing rate. They demonstrate that experimental observations fall within the feature distribution generated by the simulations. The top right panel shows a two-dimensional principal component representation of the full summary-statistics feature space (4 observed features × 4 summary statistics). Whereas the other panels show that the experimental observations are covered for each feature considered separately, the principal component representation shows that they are also covered when all features are considered jointly. VL: Vastus lateralis; VM: Vastus medialis; GM: Gastrocnemius medialis; SOL: Soleus; FDI: First dorsal interosseous, TA: Tibialis anterior; MVC: Maximal voluntary contraction.

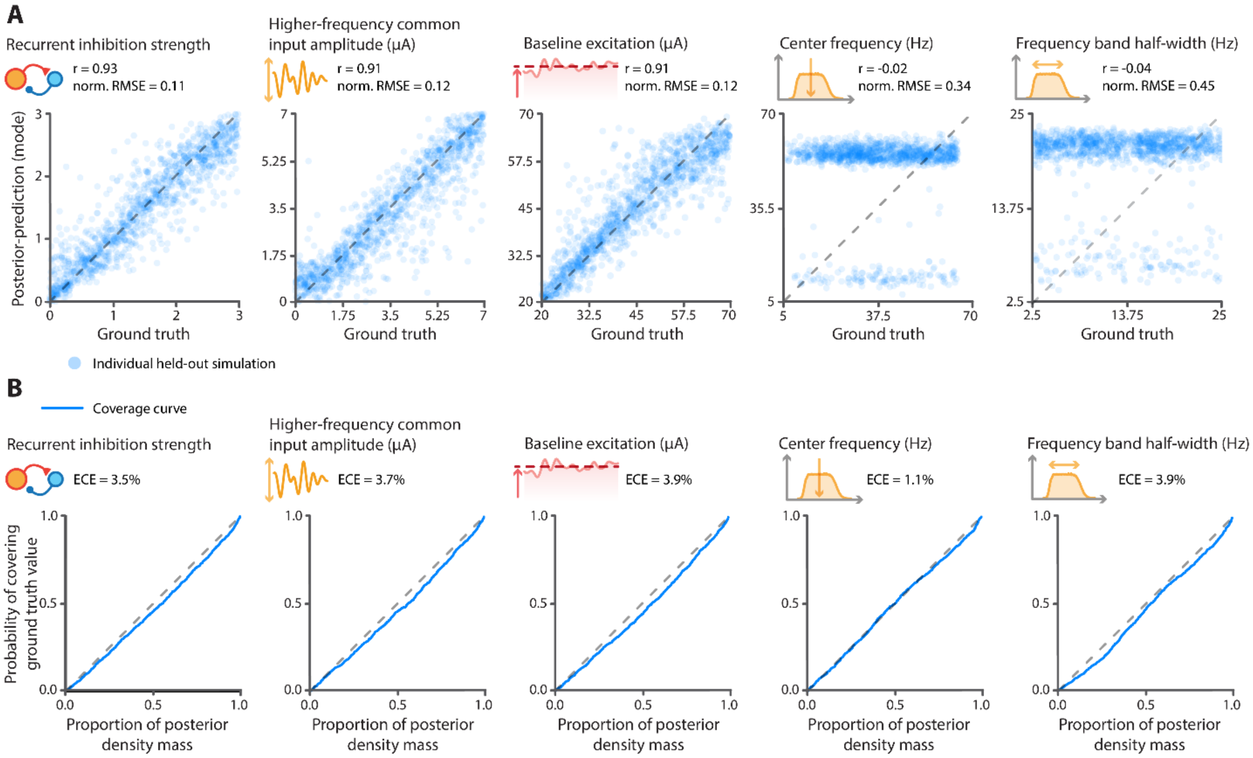


**Figure S2. Validation of inference on held-out simulated data. (A)** Accuracy of parameter recovery. The panels show, for each parameter, the relationship between the simulated ground-truth (x-axis) and the corresponding *posterior*-predicted mode (y-axis). *Norm. RMSE*: root mean square error for the [0,1]-normalized parameters. Under a uniform *prior*, expected normalized RMSE ≅ 0.41, and expected Pearson's r = 0. **(B)** Calibration of *posterior* uncertainty. Each panel shows the calibration of the neural density estimator for one parameter. The expected calibration error (ECE) is the average deviation from the identity line, i.e., perfect calibration.

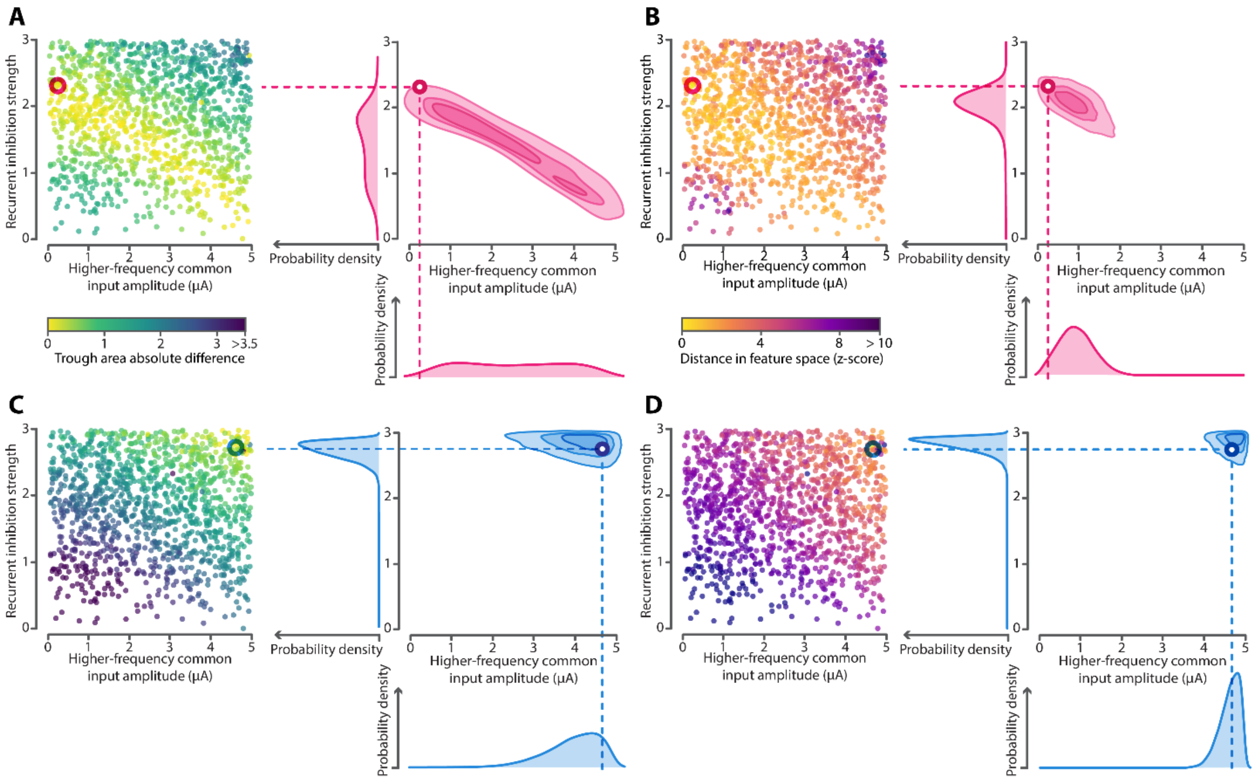


**Figure S3.** ***Posteriors* in ambiguous *vs* unambiguous simulation examples. (A)** These figures are based on the same simulated dataset as in Fig. 3A-B. Each dot represents a simulation, with color indicating the absolute difference in mean trough area relative to the selected simulation example (red circle). The right panel shows the corresponding joint *posterior* when inference is conditioned on the mean trough area alone. The resulting broad ridge-shaped *posterior* indicates strong indeterminacy. **(B)** The same data as in panel A, but colors now indicate the z-score–normalized Euclidean distance to the selected simulation (same as in Panel A) in the full summary-statistics feature space (4 observed features × 4 summary statistics), and inference is conditioned on this full feature set. The resulting *posterior* is more constrained than in panel A. **(C)** Same as panel A, but for a second selected simulation example located in a less ambiguous region of the parameter space. **(D)** Same as panel B, but for the simulation example shown in C. For this second example, uncertainty is already reduced in panel C and becomes very low in panel D.

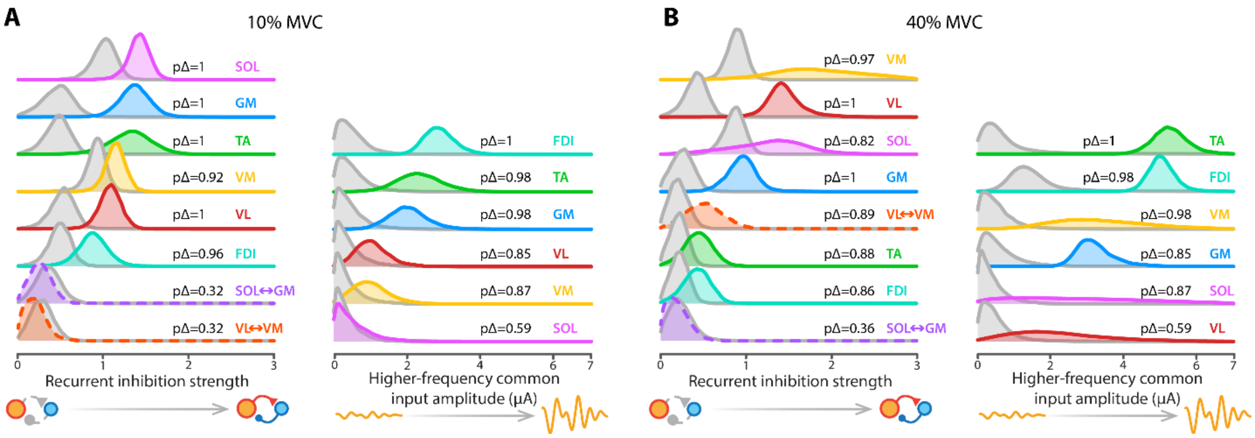


**Figure S4. Comparison of *posteriors* inferred from original and shuffled experimental data.** **(A)** Marginal *posterior* distributions of recurrent inhibition strength and higher-frequency common input amplitude are shown for each muscle and muscle pair at 10% MVC. Colored distributions correspond to inference from the original experimental data whereas gray distributions were obtained after independently circularly shifting each motor unit spike train. pΔ values indicate the probability that the parameter inferred from the original data exceeds that inferred from the shuffled data. **(B)** Same as in panel A for 40% MVC. Note that slight differences between the *posteriors* shown here and those shown in Fig. 6 are due to variability in *posterior* sampling from the same trained neural density estimator. VL: Vastus lateralis; VM: Vastus medialis; GM: Gastrocnemius medialis; SOL: Soleus; FDI: First dorsal interosseous, TA: Tibialis anterior; MVC: Maximal voluntary contraction.

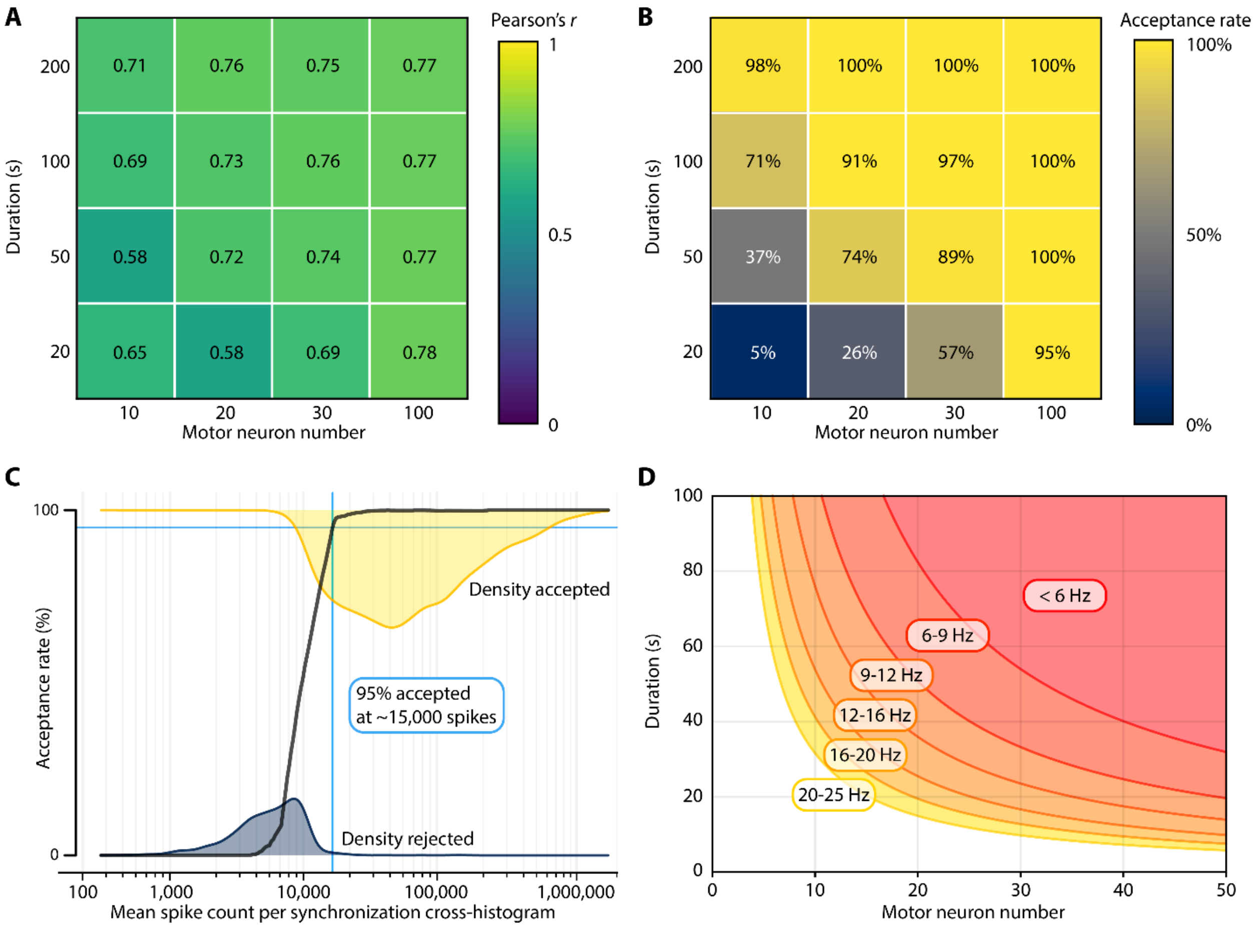


**Figure S5. Sensitivity of the simulation-based inference pipeline to recording duration, motor neuron number, and spike count. (A)** Mean Pearson's *r* between ground-truth recurrent inhibition strength and its inferred value (*posterior* mode) across simulations is shown for different recording durations and numbers of sampled motor neurons. Only simulations that passed the quality-control criteria (see Methods) are included. **(B)** Percentage of simulations retained for inference across different recording durations and numbers of sampled motor neurons. **(C)** Relationship between the mean spike count per synchronization cross-histogram and the acceptance rate for inference. Yellow and dark blue shaded areas show the densities of accepted and rejected simulations, respectively, and the black curve shows the corresponding acceptance rate. In this simulation set, an average of ~15,000 spikes per synchronization cross-histogram was associated with a 95% acceptance rate, i.e. a 95% probability that the quality-control criteria are met. **(D)** Practical guideline derived from the same simulations, relating recording duration (y-axis), motor neuron number (x-axis) and mean firing rate (color) associated with a ≥95% probability of passing the quality-control criteria.
